# Dissecting the RNA binding capacity of the multi-RRM protein Rrm4 essential for endosomal mRNA transport

**DOI:** 10.1101/2025.02.12.636894

**Authors:** Nina Kim Stoffel, Srimeenakshi Sankaranarayanan, Kira Müntjes, Anke Busch, Julian König, Kathi Zarnack, Michael Feldbrügge

**Author notes:** Co-corresponding authors: Prof. Dr. Kathi Zarnack, Theodor Boveri Institute for Biosciences, Biocenter, Julius Maximillians University Würzburg, Am Hubland, 97074 Würzburg, Germany; Prof. Dr. Michael Feldbrügge, Institute of Microbiology, Collaborative Research Center Microbial Networking Heinrich Heine University Düsseldorf, 40204 Düsseldorf, Germany. Shared first authors (Nina Kim Stoffel, Srimeenakshi Sankaranarayanan).

## Abstract

RNA-binding proteins (RBPs) utilize multiple RNA-binding domains (RBDs) to engage with extensive mRNA networks. Understanding the intricate interplay of modular RBDs is essential for uncovering RBP function. Yet, how individual RBDs shape transcriptome-wide interactions remains poorly understood. Here, we dissect the roles of the three RNA recognition motifs (RRMs) in the endosomal mRNA transporter Rrm4 during polar growth of *Ustilago maydis*. Using a comparative mutant-based iCLIP2 approach, we disclose an extensive inventory of RRM-specific binding sites. Most binding sites are prominently governed by RRM3, however, they are not critical for function. Conversely, functionally essential binding sites are recognized by a more complex RBD interplay, involving RRM1 and/or RRM2 with partial support from RRM3. By integrating transcriptome-wide RNA binding data with transcriptomics, we pinpoint their function as regulatory RNA elements affecting mRNA abundance, linking endosomal transport to stability. The modular RNA binding of Rrm4 defines distinct RNA regulons controlling mitochondrial activity, polarity factors, and cell wall remodeling – processes critical for polar growth. These findings disclose the intricate binding modes of an RBP *in vivo*, emphasizing how multiple RBDs differentiate functional binding sites from accessory ones to determine mRNA fate.

**Graphical abstract:** Figure 1.

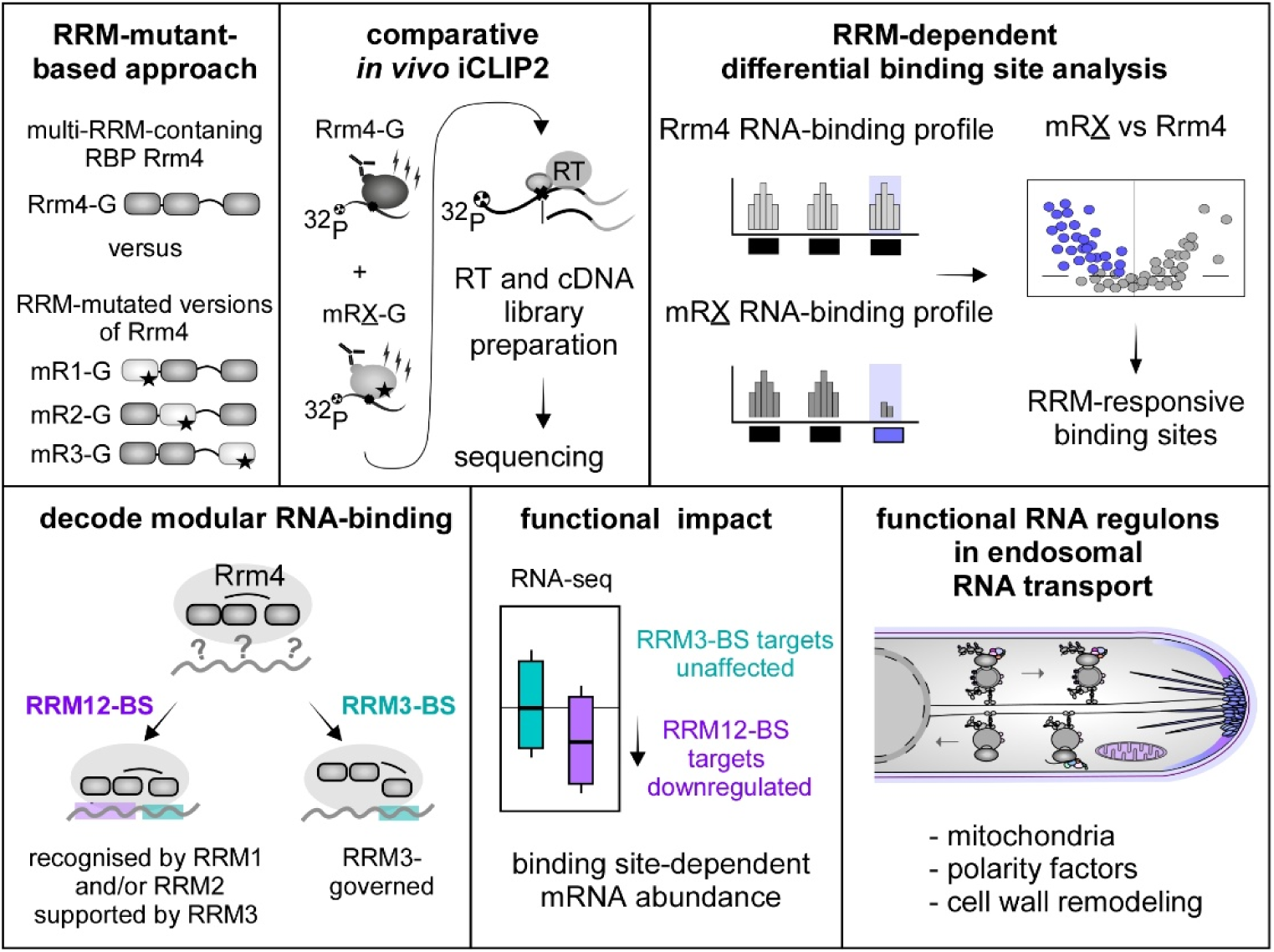

## Introduction

RNA-binding proteins (RBPs) dynamically regulate the life cycle of each mRNA from synthesis to decay. Their malfunction can lead to death and disease (1,2). Consequently, RBPs have garnered significant interest in drug development and cancer treatment (3–5). Powerful approaches to study the RNA interactome of RBPs include transcriptome-wide UV crosslinking-based methods, such as individual-nucleotide resolution UV crosslinking and immunoprecipitation (iCLIP) and its derivatives (6). In recent years, large-scale binding maps of 150 human RBPs revealed thousands of RNA binding sites (7,8), yet discerning functional binding sites from others within the full spectrum of RNA interactions presents a significant challenge. This is further complicated by the modularity of RNA-binding domains (RBDs) in most RBPs, which often mediates target recognition by binding specific RNA sequences or structures (9–11). Dissecting the intricate RNA binding code is crucial to elucidate the precise role of each RBD in recognizing and modulating functionally critical binding sites.

The RNA recognition motif (RRM) is the most prevalent RBD in eukaryotes with a characteristic β1-α1-β2-β3-α2-β4 topology (Fig. EV1A) (12,13). Canonical RNA contact regions (RNP1 and RNP2) within the β-strands are crucial for sequence-specific binding (Fig. EV1A). However, single domains are often insufficient to confer specificity and affinity (14). Hence, multiple RBDs frequently occur within RBPs enhancing RNA binding (13). Tandem RRMs, for example, provide an expanded RNA-binding surface, promoting binding specificity and avidity (15). A typical example is the ELAV family of multi-RRM RBPs (embryonic lethal abnormal vision, Fig. EV1A), which feature a typical arrangement of tandem RRMs separate from a third RRM (16). Members of this family include proteins such as HuD (ELAV-like protein 4, *Homo sapiens*), Rbp9 (*Drosophila melanogaster*), and Rrm4 (*Ustilago maydis*) regulating translation, stability, and transport of mRNAs (Fig. EV1A) (17–19). HuD, for example, is specifically implicated in neurogenesis in motor neuron function, as knock-out mice exhibit motor deficits and its loss leads to increased self-renewal of neuronal stem cells and progenitors (20,21).

We investigate the long-distance transport along microtubules, focusing on the co-transport of mRNAs on the cytoplasmic surface of early endosomes. This vesicle hitchhiking process was initially discovered in fungi but has recently been observed in other systems, including plants, mammalian cells, and neurons (22–27). In the fungal microorganism *U. maydis*, the highly polarized infectious hyphae rely on the key endosomal mRNA transport protein Rrm4 (28–30). Besides its three N-terminal RRMs, Rrm4 harbors three C-terminal MademoiseLLE domains that serve as a binding platform for interaction with the endosomal anchor protein Upa1 (Fig. 1A, Fig. EV1A) (31). Additionally, Upa2 functions as a scaffold protein, stabilizing higher-order messenger ribonucleoprotein particle (mRNP) complexes, including the RNA chaperone Grp1 and the poly(A)-binding protein Pab1 (Fig. EV1B) (32).

**Figure 1.**
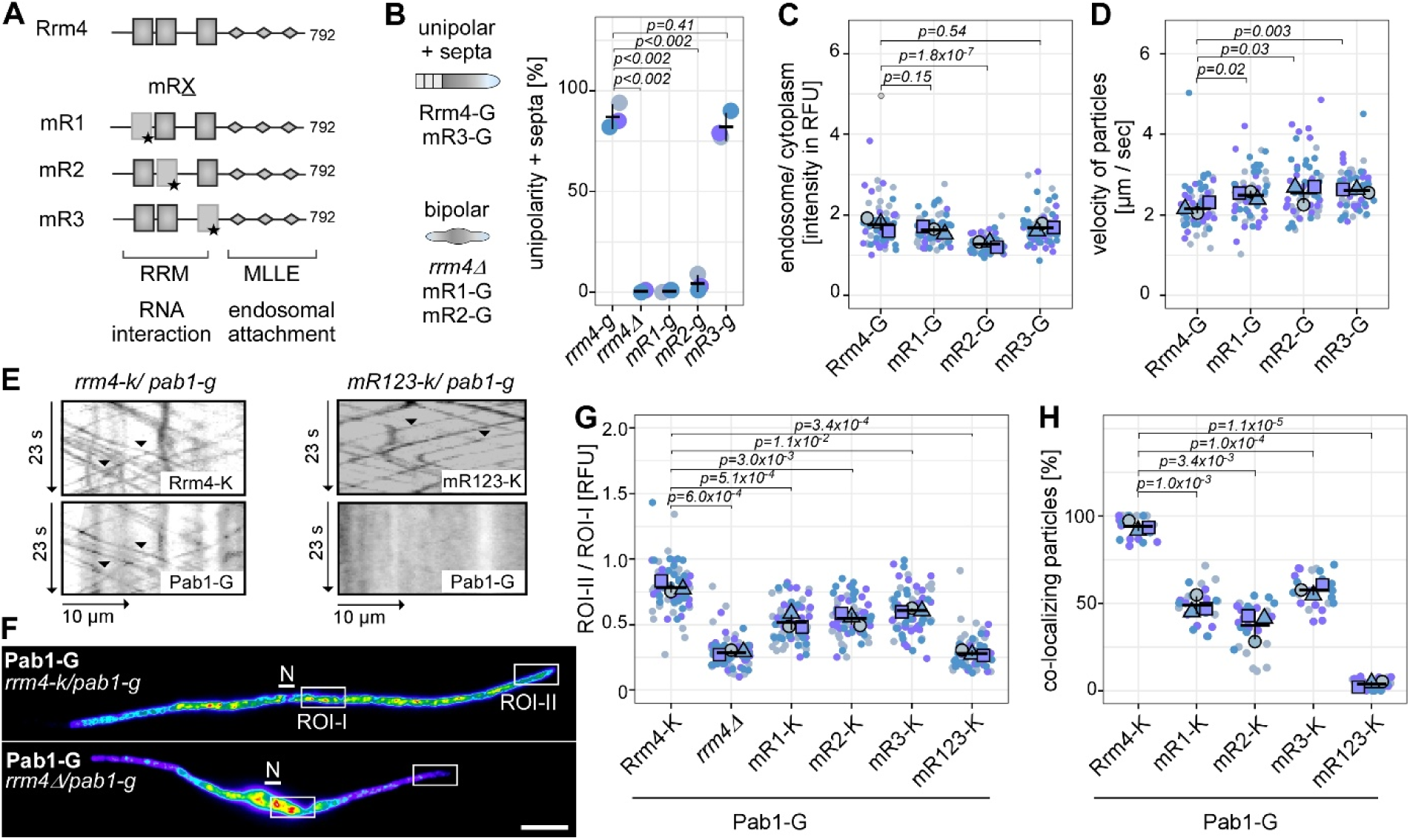
**Mutations in RRM domains cause differential defects during polar growth.** (**A**) Schematic representation of Rrm4 variants (792 amino acids): grey rectangles, RNA recognition motif (star indicate mutation); diamonds, MLLE domains. (**B**) Hyphal growth (6 h. p. i.) Unipolar hyphae with inserted septa were quantified (Student’s *t*-test, unpaired; three biological replicates are shown in different blue shadings, N > 100) (**C**) Ratio of Gfp relative fluorescence unit (RFU) intensity comparing endosomes versus cytoplasm. Blue shading indicates results from three independent experiments (Student’s *t*-test, unpaired; based on the means of the three biological replicates). The mean of each replicate is given in different forms and respective colors (replicate 1=circle; replicate 2=triangle, replicate 3=square; n > 60). (**D**) Velocity of Rrm4-containing particles. Shading indicates results from three independent experiments (Student’s *t*-test, unpaired; three biological replicates, N > 60). (**E**) Kymographs (inverted fluorescence images) of hyphae (6 h. p. i.) showing mKate2 (-K) and Gfp (-G) signals at the top and bottom. (**F**) False color imaging of Pab1-G fluorescence in hyphae (6 h. p. i.). Regions of interest (ROI) are marked by rectangles (size bar, 10 µm). (**G**) Quantification of Pab1-G distribution comparing ROI-I (perinuclear) versus ROI-II (growth pole). Shading indicates results from three independent experiments (Student’s *t*-test, unpaired; N > 60). (**H**) Quantification of Rrm4-K versions co-localising with Pab1-G-containing particles (Student’s *t*- test, unpaired; three biological replicates, N > 30).

Loss of Rrm4 results in aberrant hyphal polarity, linking endosomal mRNA transport to unipolar growth (Fig. 1B, Fig. EV1D-E) (33). One function of endosomal transport is the widespread distribution of bulk mRNAs and associated ribosomes throughout the cell (34,35). Additionally, Rrm4 specifically targets all four septin mRNAs (*cdc3*, *cdc10*, *cdc11*, *cdc12*) and is required for their endosomal transport, a process crucial for the formation of heteromeric septin complexes and their proper localization at the cell tip, essential for defined polarity (32,36,37). Previous iCLIP analysis of Rrm4 identified hundreds of mRNA targets preferentially bound in the 3’ untranslated region (UTR) (34). The motif UAUG was enriched in Rrm4 target mRNAs, and yeast-3-hybrid assays suggested that the third RRM (RRM3) is essential for binding UAUG (34). However, mutational analyses indicated that RRM3 is dispensable for unipolarity, while RRM1 is crucial, indicating that each RRM contributes differently to the role of Rrm4 in polar growth (33). Here, we integrated a comprehensive iCLIP2 analysis of mutated Rrm4 variants with transcriptomics and live-cell imaging to decode the modular RRM recognition code and unveil its functional significance.

## Material and methods

### *U. maydis* strain generation, cultivation of yeast cells and induction of hyphal growth

The laboratory strain AB33 of *U. maydis* (homotypic synonym of *Mycosarcoma maydis*, NCBI taxonomy ID 5270) was used to synchronize hyphal growth by altering the nitrogen source. In brief, cell suspensions were grown in a complete medium containing 1% glucose at 28°C with constant agitation at 200 rpm until an OD_600_ of 0.5 was reached. Hyphal growth was induced by transferring the cells to a nitrate minimal medium supplemented with 1% glucose and incubating them at 28°C with agitation at 200 rpm for 6 hours post-induction (h. p. i.) (38). Further details regarding media composition, growth conditions, and general strategies for strain generation in *U. maydis* were previously described (38,39). Strains, plasmids and DNA oligonucleotides used for strain generation are given in the supplementary information (Supplementary Tables S1-S3). UMAG identifiers for selected Rrm4 target genes are given in Supplementary Table S4. Strain names are typically written in italics and are designated based on the construct they carry (*rrm4-g*, *mR1-g*, *mR2-g*, *mR3-g*) or, in cases where specific genes are absent, marked with a delta symbol (*rrm4Δ*). For example, *rrm4-g* refers to a strain expressing the Rrm4-G protein. Protein names are written without additional tags when referring to them in a general sense.

### Microscopy

For standard microscopy, cells were grown in 20 mL cultures to an OD_600_ of 0.5. Hyphal growth was induced and cells were imaged 6 h. p. i. To assess unipolar and bipolar growth, more than 100 hyphae (6 h. p. i.) per strain were analyzed across three biological replicates (n=3). To quantify the number of shuttling particles, their velocity, and the endosomal attachment of fluorescently labeled fusion proteins (Gfp, mKate2, Rfp, mCherry), movies were recorded with an exposure time of 150 ms for a total of 150 frames (32). For generating kymographs, 15 or 30 μm sections of hyphal cells starting from the cell tip or perinuclear region were used. Only processively moving particles that traveled >5 μm were considered shuttling particles. Each experiment involved more than 24 hyphae per strain (n=3). Rrm4-G signal intensity of shuttling particles was measured using kymograph intensity relative to the cytoplasmic background (relative fluorescent units, RFU). All movies, kymographs, and images were processed and analyzed using Metamorph software (Version 7.7.0.0, Molecular Devices, Seattle, IL, USA). For co-localization experiments of shuttling particles, a two-channel imaging system (DV2; Photometrics, Tucson, AZ, USA) was utilized. Co-localization efficiency was assessed by calculating the ratio of Pab1-Gfp particles to Rrm4-mKate2 particles. Data analysis and visualization were conducted in RStudio (version R 4.4.2).

### Western blot analysis

Protein concentrations of cell extracts were measured using either the BCA assay (Thermo Fisher, Pierce™ 23225) or the Bradford assay (BioRad). Samples were adjusted to 1 mg/mL in NuPAGE LDS containing DTT according to the manufacturer’s instructions (Thermo Fisher, NP0007). Samples were boiled for 5 minutes and loaded onto a 4-12% NuPAGE™ Bis-Tris gel (Thermo Fisher, NP0321). Electrophoresis was carried out at 180 V for 1.5 hours. Proteins were transferred onto a nitrocellulose membrane (Amersham Protran) via wet blotting using a NuPAGE™ transfer buffer supplemented with 15% methanol (Thermo Fisher, NP00061) for 1.5 hours at 30 V. For Western blot analysis, primary antibodies α-GFP (1:2000 dilution; Roche, 11814460001) and α-Actin (1:2000; MP Biomedicals, 0869100) were applied for 1 hour at room temperature under continuous rotation. Subsequently, the membrane was washed three times with TBS-T for 15 minutes each. The secondary antibody, α-mouse HRP conjugate (1:5000 dilution; Promega, W4021), was applied for 1 hour. All antibodies were diluted in TBS-T containing 5% milk powder. The detection was performed using LAS400 (GE Healthcare) and ECL™ Prime (Amersham, RPN2232) following the manufacturer’s guidelines.

### Comparative microbial iCLIP2

The comparative iCLIP2 approach used 150 mL hyphae cultures (OD_600_ =0.5, 6 h. p. i.) as the starting material. All experiments were conducted as detailed in the microbial iCLIP2 protocol tailored for Rrm4-G and mutated versions (40). Rrm4-G and the comparative mutants (mR1- G, mR2-G, mR3-G) were prepared simultaneously in at least five biological replicates for high-resolution analysis. The iCLIP2 libraries were sequenced in two independent runs (1: Rrm4-G, mR1-G; 2: Rrm4-G, mR2-G, and mR3-G) to obtain at least 25 million reads per sample using an Illumina NextSeq 500 platform to produce 92 nt single-end reads. These reads included a 6 nt sample barcode and unique molecular identifiers (UMIs, Supplementary Table S3) of 5+4 nt (41).

### Processing of iCLIP reads

The processing of iCLIP2 reads as well as the binding site definition (see below) were carried out mostly as previously described (42). In brief, FastQC (version 0.11.8; http://www.bioinformatics.babraham.ac.uk/projects/fastqc/) was utilized for initial quality controls. Reads with a Phred quality score below 10 in the barcode or UMI regions were removed from further analysis using the FASTX-Toolkit (version 0.0.14; https://github.com/agordon/fastx_toolkit) and seqtk (version 1.3; https://github.com/lh3/seqtk/). Flexbar (version 3.4.0) (43) was used to de-multiplex the reads based on the sample barcode on positions 6 to 11 and to trim the barcodes, UMI regions, and adapter sequences from the ends of the reads, while ensuring a minimal overlap of 1 nt between the read and the adapter. UMIs were added to the read names and only reads with a minimum length of 15 nt were retained. Trimmed reads were mapped to the *U. maydis* genome and its annotations (Ensembl release 51; assembly version Umaydis521_2.0) (44) using STAR (version 2.7.3a) (45). During mapping, up to 4% mismatched bases were accepted and soft-clipping was prohibited on the 5’ end. Reads directly mapped to the chromosome ends were removed since they do not have an upstream position and, consequently, no crosslink position can be extracted. PCR duplicates were removed from uniquely mapping reads using UMI-tools (version 1.0.0) (46). Crosslinked nucleotides were extracted using the BEDTools suite (version 2.27.1) (47) as described in Chapter 4.2 of (42).

### iCLIP2 binding site definition

Peak calling was performed using PureCLIP (version 1.3.1) (48) with default parameters on the merged, preprocessed alignments of the individual replicates. The resulting crosslink sites were used to define binding sites with the R/Bioconductor package BindingSiteFinder (version 1.7.9) (42); (https://doi.org/doi:10.18129/B9.bioc.BindingSiteFinder) (49). Focusing on the most significant sites, we excluded the lowest 5% of crosslink sites ranked by the PureCLIP score. Additionally, a gene-wise filter was applied to remove the bottom 10% of crosslink sites based on PureCLIP scores for each gene. Analysis of crosslink event coverage revealed that binding sites of width 5 nt more effectively captured the central peak in iCLIP2 read coverage than the previously used 9 nt width. Consequently, crosslink sites located within a 5 nt distance were merged into binding regions. Isolated crosslink sites and binding sites of width 2 nt were removed. Binding site centres were defined at crosslink events with the highest PureCLIP scores and extended by 2 nt on either side. For binding sites longer than 5 nt, the binding regions were divided into 5 nt regions by iteratively identifying the maximum signal and extending by 2 nt on either side, ensuring no overlap between binding regions. Reproducibility was assessed by retaining binding sites that contained more crosslink events than the 5th percentile of the crosslink distribution across all binding sites, with a minimum of two crosslink events per site. Sites were considered reproducible if detected in at least n−1 replicates, where n is the total number of replicates. To assign gene and transcript regions to the binding sites, we utilized the gene annotation file version *U. maydis* 521_2.0.41 (http://ftp.ensemblgenomes.org/pub/fungi/release-53/fasta/ustilago_maydis/dna/) (50). Genes in the *U. maydis* genome were manually extended by 300 nt on each side to encompass potential 5’ UTR and 3’ UTR regions (18). To resolve overlapping transcript types, we applied the following hierarchy: tRNA > mRNA > rRNA > snRNA > snoRNA. For overlapping transcript regions, binding sites were assigned to the transcript regions (5’ UTR, CDS, intron, 3’ UTR) that were most frequently observed.

### Differential binding analysis

The differential binding analysis was performed using the package BindingSiteFinder (version 1.7.9) (51). The binding sites for each iCLIP2 dataset were defined independently as mentioned above. For the differential binding analysis, the binding sites from the Rrm4 iCLIP2 data were used as control. Binding sites from all datasets (Rrm4-G, mR1-G, mR2-G, and mR3-G iCLIP2 data) were combined to enable differential testing. For pairwise comparisons, the binding sites corresponding to the Rrm4 control and the corresponding mutant condition were extracted from the merged dataset. Differential testing was performed by dividing crosslink events for each gene into two categories: the binding site bins, containing crosslink events overlapping the binding sites, and the background bin, encompassing all crosslink events within the gene that do not overlap with binding sites or the adjacent 2 nt offset regions flanking the binding sites. The inclusion of the offset regions ensures that any signal near the binding sites is excluded from the background bin, allowing for a more accurate representation of the background bin. Additionally, the background bin accounts for variations in RNA expression levels. Subsequently, the crosslink counts for each binding site and the corresponding background bin were extracted for each gene. The background bin counts were used to normalize and correct for expression level changes. Genes were excluded from the analysis if they met any of the following criteria: fewer than 100 crosslink events across all samples, more than 30% of crosslink events attributed to binding sites, more than 70% of crosslink events attributed to the background, or more than 95% of total crosslink events attributed to a single condition. Finally, fold change for each binding site between the two conditions was calculated using the DESeq2 likelihood ratio test (52), separating transcriptional changes from alterations at the binding site level.

### RNA isolation and sequencing

25 mL of hyphal cell culture with an OD_600_ of 0.5 was used to isolate total RNA. Cells were harvested by centrifugation (7,150 *g*, 10 min, 4°C) and resuspended in 2 mL of PBS (phosphate buffered saline) divided and transferred into 2 mL reaction tubes. Cells were harvested by another round of centrifugation (16,200 *g*, 5 min, 4°C) and tubes containing the cell pellets were frozen in liquid nitrogen. A steel bead (5 mm) was added to each cell pellet and placed in the pre-cooled TissueLyser Adapter (Qiagen). The samples underwent two rounds of mechanical disruption for 30 seconds at 30 Hz each (Retsch, M400), with a cooling step in liquid nitrogen between rounds. The cryogenically frozen cell powder was resuspended in 500 µL of TriZol (Invitrogen) and incubated, shaking at room temperature for 5 minutes at 1000 rpm. Next, 500 µL of RNase-free water was added, and the mixture was manually shaken for 30 seconds. Subsequently, 400 μL of chloroform/isoamylalcohol (24:1; BioChemica) was added. The samples were shaken vigorously by hand for 3 minutes, then incubated at room temperature for another 3 minutes. Following this, the samples were centrifuged for 10 minutes at 16,200 g. The aqueous phase was transferred to a fresh 1.5 mL tube pre-filled with 250 μL of chloroform/isoamylalcohol (24:1; BioChemica). This extraction process was repeated, followed by centrifugation at 16,200 g for 5 minutes at 4°C. The aqueous phase was transferred to a tube containing 40 μL of sodium acetate (3 M, pH 5.2) and 1 mL of ice-cold 100% ethanol. Samples were incubated at -20°C for 2 hours, and RNA was pelleted by centrifugation at 16,200 g for 30 minutes at 4°C. The pellet was washed twice with 75% ice-cold ethanol, centrifuged for 5 minutes at 16,200 g and 4°C, briefly dried, resuspended in 50 μL of RNase-free water, and stored at -80°C. RNA concentration was quantified using a NanoDrop spectrophotometer (Thermo Fisher Scientific). For each sample, 2 µg of RNA was subjected to DNase digestion using the dsDNase Kit (Thermo Fisher, EN0771) at 37°C for 10 minutes. The RNA was then precipitated overnight by adding 1 mL of ice-cold 100% ethanol and 40 μL of sodium acetate (3 M, pH 5.2). The RNA pellet was collected by centrifugation at 16,200 g for 30 minutes at 4°C, washed twice with 75% ice-cold ethanol and centrifuged again for 5 minutes at 16,200 g and 4°C. The pellet was briefly dried, resuspended in 10 μL of RNase-free water, and stored at -80°C. RNA concentration was quantified using the Qubit RNA BR Assay (Invitrogen by Thermo Fisher Scientific), and RNA integrity was verified with the Bioanalyzer RNA Nano Assay (Agilent). NGS library prep was performed with Illumina’s Stranded mRNA Prep Ligation Kit following Stranded mRNA Prep Ligation ReferenceGuide (April 2021, Document 1000000124518 v02). Libraries were prepared with a starting amount of 500 ng and amplified in 11 PCR cycles. Two post-PCR purification steps were performed to exclude residual primer and adapter dimers. Libraries were profiled in a High Sensitivity DNA chip on a 2100 Bioanalyzer (Agilent Technologies) and quantified using the Qubit dsDNA HS Assay Kit, in a Qubit 4.0 Fluorometer (Invitrogen by Thermo Fisher Scientific). All samples were pooled together in equimolar ratio aiming for at least 10 million reads per sample and sequenced on a NextSeq 2000 P3 (100 cycles) FC, SR for 1x116 cycles plus 2x10 cycles for the index reads and one dark cycle upfront R1.

### Differential gene expression analysis

Basic quality control of RNA-seq libraries was conducted using FastQC (version 0.12.1; https://www.bioinformatics.babraham.ac.uk/projects/fastqc/). Prior to mapping, RNA-seq reads were processed using Trimmomatic (version 0.39) (53) to remove adapter sequences and enhance sequence quality. Trimming was performed under the following criteria: Reads were truncated when the average Phred score within a 4 nt sliding window fell below 15, reads were trimmed at the start or end of reads when the Phred score dropped below 30 and reads shorter than 36 nt were discarded. The processed reads were aligned to the *U. maydis* genome using STAR (version 2.7.11b) (45), allowing up to 4% mismatches. Reads per gene were counted with featureCounts (version 2.0.1) (54) using default parameters. Differential gene expression analysis between the wild-type (*rrm4-g*) and mutant conditions (*rrm4Δ, mR1-g*, *mR2-g*, *mR3-g* and *mR123-g;* Supplementary Table S1) were performed using DESeq2 (version 1.42.0) (52) in the R environment (version 4.3.1; https://www.R-project.org/).

### Motif analysis

*De novo* motif discovery analysis was conducted using the XSTREME tool from the MEME suite (55). A 15 nt window encompassing binding sites extended by 5 nt to either side was used as input. Shuffled input sequences served as a background control. XSTREME analysis was conducted with default parameters. For each enriched motif, its relative enrichment ratio was compared against the absolute percentage of binding sites with this motif. The motif logo was created using the package ggseqlogo (version 0.2) (56) in R. For *k*-mer analysis, the frequency of all possible pentamers within the same 15 nt windows was computed using the R/Bioconductor package Biostrings (version 2.70.1, https://bioconductor.org/packages/ Biostrings.). Additionally, pentamer frequency was calculated within randomly selected 15 nt regions from the genome, which served as a control. Motif enrichment was determined as the ratio of pentamer frequency in binding sites to its total frequency across both the binding sites and background regions. Since the antisense and scrambled sequences of UAUG contain parts of the motif, we chose the random GCAU motif as a control.

### Positional analysis of binding sites

RRM12-BS and RRM3-BS were each extended by 150 nt from their centres to obtain a 301 nt window. For relative positional analysis, the overlaps of RRM12-BS within the 301 nt RRM3- BS window, and *vice versa*, were identified. The relative position of RRM12-BS within the RRM3-BS window (or RRM3-BS within the RRM12-BS window) was determined by calculating the distance from the start position of the binding site to the centre of the window.

### GO term analysis

For the functional enrichment analysis, we stratified the mRNA targets (RRM12-BS or RRM3- BS targets) into groups based on the number of binding sites they possess. The mRNA targets were initially defined as those containing at least one binding site ([1,>1], with subsequent groups defined as progressively more stringent [2, >2] and [3, >3]). Gene Ontology (GO) term analysis was conducted using the R package gProfiler2 (version 0.2.3) (57), applying a significance threshold at *P* value ≤ 0.05 and the g_SCS correction method. The results of the GO term analysis were visualized using heatmaps generated with the R/Bioconductor package ComplexHeatmap (version 2.18.0) (3,58,59).

### Gene set enrichment analysis (GSEA)

The GSEA analysis was performed using the R/Bioconductor package ClusterProfiler (v 4.10.1) (60). For input data, the mRNA targets of RRM12-BS and RRM3-BS were ranked by their mRNA expression levels (log_2_ fold change) in the *rrm4Δ vs. rrm4-g* RNA-seq data in descending order. Ensembl annotation for GO term Cellular Component was retrieved from the web server gProfiler (https://biit.cs.ut.ee/gprofiler/gost) (57) and was used as pre-defined gene sets. GSEA was performed with an adjusted *P* value cutoff of < 0.05, applying the Benjamini-Hochberg correction. The distribution of statistically significant gene sets in the pre-ranked target mRNA list was visualized using the R/Bioconductor package enrichplot (version 1.22.0, DOI: 10.18129/B9.bioc.enrichplot).

### Structural prediction of RRM3-UAUG

The structural modeling was performed using AlphaFold3 (3) with default settings, employing the full-length Rrm4 (A0A0D1DWZ5) and the RNA motif sequence [5’-UAUG-3’]. PyMOL (The PyMOL Molecular Graphics System, version 2.1) was utilized for visualization, focusing on the RRM3-UAUG interaction, which was vividly highlighted to enhance clarity.

### RT-PCR

Total RNA was isolated as previously described. RNA was reverse transcribed using the Maxima H Minus First Strand cDNA Synthesis Kit (Thermo Fisher), following the manufacturer’s instructions. The resulting cDNA was diluted 1:10 and used for standard PCR with the GoTaq Green Master Mix (Promega). The primers used for PCR are listed in Supplementary Table S3.

### Protein sequence comparison

To identify mammalian homologs of FERRY-associated targets (22), protein sequence comparisons were performed using the BLASTp program (https://blast.ncbi.nlm.nih.gov/Blast.cgi) (61). The analysis aimed to identify *U. maydis* orthologs by querying the non-redundant protein sequences (nr) database. The default parameters used included an E-value threshold of 10, the BLOSUM62 substitution matrix, a word size of 6, and gap costs set to 11 for existence and 1 for extension. The analysis was conducted bidirectionally, from human to *U. maydis* and *vice versa* (Supplementary Table S5).

## Results

### The three RRM domains differentially impact hyphal growth and mRNA localization

To dissect the function of the three individual RRM domains of Rrm4 during unipolar growth, we exploited Rrm4 versions carrying four amino acid mutations in the respective RNP1 regions of the three RRMs (Fig. 1A; Fig. EV1A; mR1-3-G) (33). We generated strains in which the *rrm4* gene was replaced with variants encoding the mutated Rrm4 versions, each C-terminally fused to Gfp (e.g., mR1-G; Fig. 1A; Fig. EV1A; Supplementary Tables S1-2). Importantly, neither mRNA nor protein amounts were altered due to the mutations (Fig. EV1C).

Analyzing hyphal growth revealed that strains expressing mR1-G and mR2-G, with mutations in RRM1 and RRM2, respectively, resembled the *rrm4* deletion (*rrm4Δ*) strain, exhibiting an increased amount of bipolarly growing cells (Fig. 1B; Fig. EV1D-E). Both mutant versions, however, are still shuttled on endosomes (Fig. EV1F). The RRM2 mutation differed from the RRM1 mutation, as the rate of bipolar hyphae with septa was increased (Fig. EV1E), while the ratio of endosome-associated-to-cytoplasmic mR2-G intensity was decreased, indicating less efficient attachment (Fig. 1C). By contrast, mutations in RRM3 (mR3-G) did not significantly affect the rate of unipolar growth, the number of shuttling particles nor the signal intensity of Rrm4 on shuttling endosomes (Fig. 1B-C; Fig. EV1D-F) (33). Interestingly, all three RRM mutations increased the velocity of endosomes (Fig. 1D). A similar effect was observed in the mutant lacking the scaffold protein Upa2 (*upa2Δ*), suggesting elevated motility due to the reduced burden of transporting mRNA ribonucleoprotein (mRNP) cargo and associated ribosomes (32).

To study mRNP trafficking, we used the poly(A)-binding protein Pab1-G as a reporter for bulk mRNA transport (Fig. 1E-F; functional C-terminal eGfp fusion) (36). Pab1 is associated with the poly(A) tail of most mRNAs (62,63) and its visualization can hence be used as a proxy for the localization of mRNAs. As expected, co-localization was almost completely abolished in the mR123-K-expressing strain (C-terminal mKate2 fusion carrying mutations in all three RRM domains of Rrm4; Fig. 1E, H; Supplementary Tables S1-2). This is consistent with previous observations that mRNA association with Pab1 on endosomes is RRM-dependent (36). As expected, the co-localization of Pab1-G was reduced in strains with mutations in single RRMs (Fig. 1E, H). Comparing the subcellular localization of Pab1-G at the growth apex with the perinuclear region revealed an almost equal distribution of Pab1-G signals in Rrm4 wild-type cells (Fig. 1F-G, Fig. EV1G). However, in *rrm4Δ* or *mR123-k* strains, Pab1-G localization was reduced at the growth pole and increased in the perinuclear region (Fig. 1F-G; Fig. EV1G). Individual RRM mutations led to intermediate effects on Pab1-G localization (Fig. 1G, Fig. EV1G), suggesting that mRNA transport is still operational but to various extents.

In summary, the RRM domains play distinct roles: RRM1 and RRM2 are essential, while RRM3 is largely dispensable for unipolar growth. Although all mutants affect Pab1-associated mRNA localization, the impact of the loss of RRM3 is the least severe, highlighting the dominant roles of RRM1 and RRM2 in mRNA localization and hyphal polarity.

### A comparative iCLIP2 approach uncovers specific signatures for all three RRMs

To study the contribution of the individual RRMs to target RNA recognition, we conducted a comparative iCLIP2 analysis using Rrm4-G and its mutants (Fig. 2A). As expected, quantification of *in vivo* UV-crosslinked RNA revealed reduced binding in all three mutants compared to Rrm4-G (Fig. 2B). In line with earlier UV crosslink experiments (33), the most drastic reduction was observed for mR3-G (Fig. 2B). To ensure sufficient recovery of UV-crosslinked RNA, particularly for RRM mutants, we optimized our iCLIP2 protocol and sequenced at least five biological replicates per experiment (40,41). The binding landscapes exhibited high reproducibility among biological replicates and across datasets (Fig. EV2A). Peak calling (48) and binding site definition (42,49) yielded a total of 50,410 binding sites across all datasets. To assess the binding specificity of the RRM domains, we compared the background-normalized crosslink counts of the mutant binding sites with those of Rrm4 wild-type. Using a differential binding analysis, we identified binding sites responsive to mutating each of the three RRM domains, i.e., showing a significant reduction in binding compared to wild-type (log_2_-transformed fold change [log_2_FC] < -1, adjusted *P* value < 0.05 [Benjamini-Hochberg correction], and mean of normalized crosslink events [baseMean] > 15). These sites were strongly affected by mutation and likely directly contacted by the corresponding RRM (defined as RRM1-, RRM2-, or RRM3-responsive sites; Fig. 2C; Supplementary Table S6). RRM3 has the highest number of responsive binding sites, followed by RRM1 and RRM2 (Fig. 2C). Additionally, we observed binding sites with increased binding following RRM domain mutation (log_2_FC > 1, adjusted *P* value < 0.05, and baseMean > 15), likely indicating aberrant binding of the mutated Rrm4 variants.

**Figure 2.**
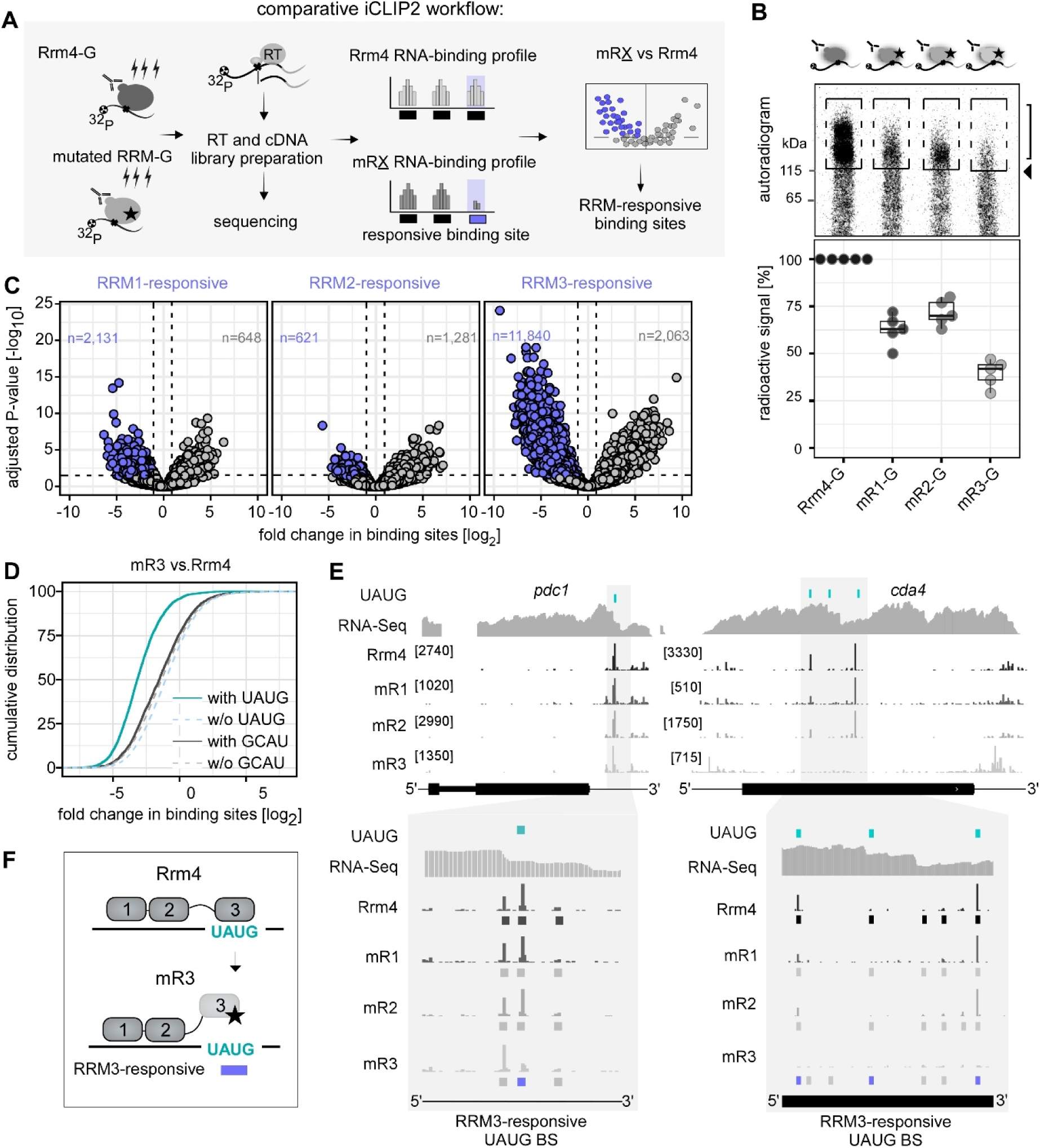
Comparative iCLIP2 approach discloses transcriptome-wide RRM3 / UAUG interaction. (**A**) Schematic representation of the comparative iCLIP2 approach. The binding landscape, derived from crosslink events in the Rrm4 mutant iCLIP2 (mRX-G=mutated RRM-G, with mutations in either RRM1, RRM2, or RRM3 domain; bottom), was compared to the wild-type Rrm4-iCLIP2 analysis (Rrm4-G; top). Binding sites dependent on the RRM domain show a significant reduction in crosslink events in the mutant compared to the wild-type (highlighted in purple). These binding sites are designated as responsive binding sites (indicated by a purple rectangle). Binding sites shown in black represent those without significant differences in crosslink events upon RRM mutation (RT, reverse transcription; G, Gfp) (**B**) Top: representative autoradiogram of UV-crosslinked radioactively labeled RNA bound to Rrm4-G or the respective mutants. Rectangles indicate regions of radioactive signal quantified at the bottom (arrowhead indicates the estimated position of full-length Rrm4-G protein). (**C**) Volcano plot showing changes in binding for mR1 (left), mR2 (center), and mR3 (right) compared to wild-type Rrm4. Binding sites exhibiting a significant reduction in binding are labeled in blue. The dotted lines indicate the significance cutoffs (absolute fold change [log_2_] > 1, adjusted P-value < 0.05, baseMean > 15). (**D**) Cumulative fraction of binding sites with (solid line) or without (dotted line) UAUG (petrol) or GCAU motifs (negative control motif; dark grey) with given binding site fold change. (**E**) Genome browser view of *pdc1* (14-3-3 gene family) and *cda4* (chitin deacetylase 4) mRNA (UAUG motif and RRM3-responsive binding sites in petrol and blue, respectively). Numbers represent the upper limit of stack heights for iCLIP2 crosslink events. (**F**) Graphical representation of Rrm4 (top) and mR3 interaction with UAUG. Mutation in RRM3 (star) cause loss of UAUG recognition.

Conducting a *de novo* motif discovery and *k*-mer analysis in the vicinity of responsive sites revealed a clear motif only for RRM3-responsive sites (Fig. EV2B-C). Here, the motif UAUG and its derivatives were most enriched (Fig. EV2B-C). More than 80% of UAUG-containing sites showed decreased binding upon RRM3 mutations compared to sites lacking the motif (Fig. 2D; EV2D). This is consistent with our previous yeast three-hybrid assays showing the interaction of RRM3 with UAUG (34). Zooming in the crosslinking landscape of mR3 reveals a clear absence of the prominent UAUG-containing sites observed in mR1, mR2, and Rrm4 (Fig. 2E). In essence, RRM3 binds the UAUG motif *in vivo* in a transcriptome-wide, sequence-specific manner (Fig. 2F) and confirms that comparative iCLIP2 enables the dissection of individual RRM binding sites.

### The tandem RRMs conjointly bind to RNA and are further supported by RRM3

Given the modular multi-RRM architecture, we analyzed whether the three RRMs influence each other’s RNA binding. While most RRM3-responsive sites were unique to the third RRM, a substantial amount of RRM1- and RRM2-responsive sites were also responsive to RRM3 (Fig. EV3A). To address this link, we compared the quantitative changes in crosslink events at all binding sites following RRM mutations (Fig. 3A). As expected, the majority of RRM3- responsive sites showed only a minimal reduction upon RRM1 and RRM2 mutation (Fig. 3A, right, compare #9, with #7 and #8). In contrast, most RRM1- and RRM2-responsive sites displayed a marked decrease in binding upon RRM3 mutation, in addition to a reduction from mutations in their respective domains (Fig. 3A, #3 and #6). Furthermore, the binding at RRM1- responsive sites decreased upon mutation in RRM2, and *vice versa* (Fig. 3A, #2 and #4), highlighting an additional interdependency within the tandem RRM domains. Thus, for a large subset of RRM1- and RRM2-responsive sites, RRM3 further supported RNA binding, and RRM1 and RRM2 also affect each other’s binding.

**Figure 3.**
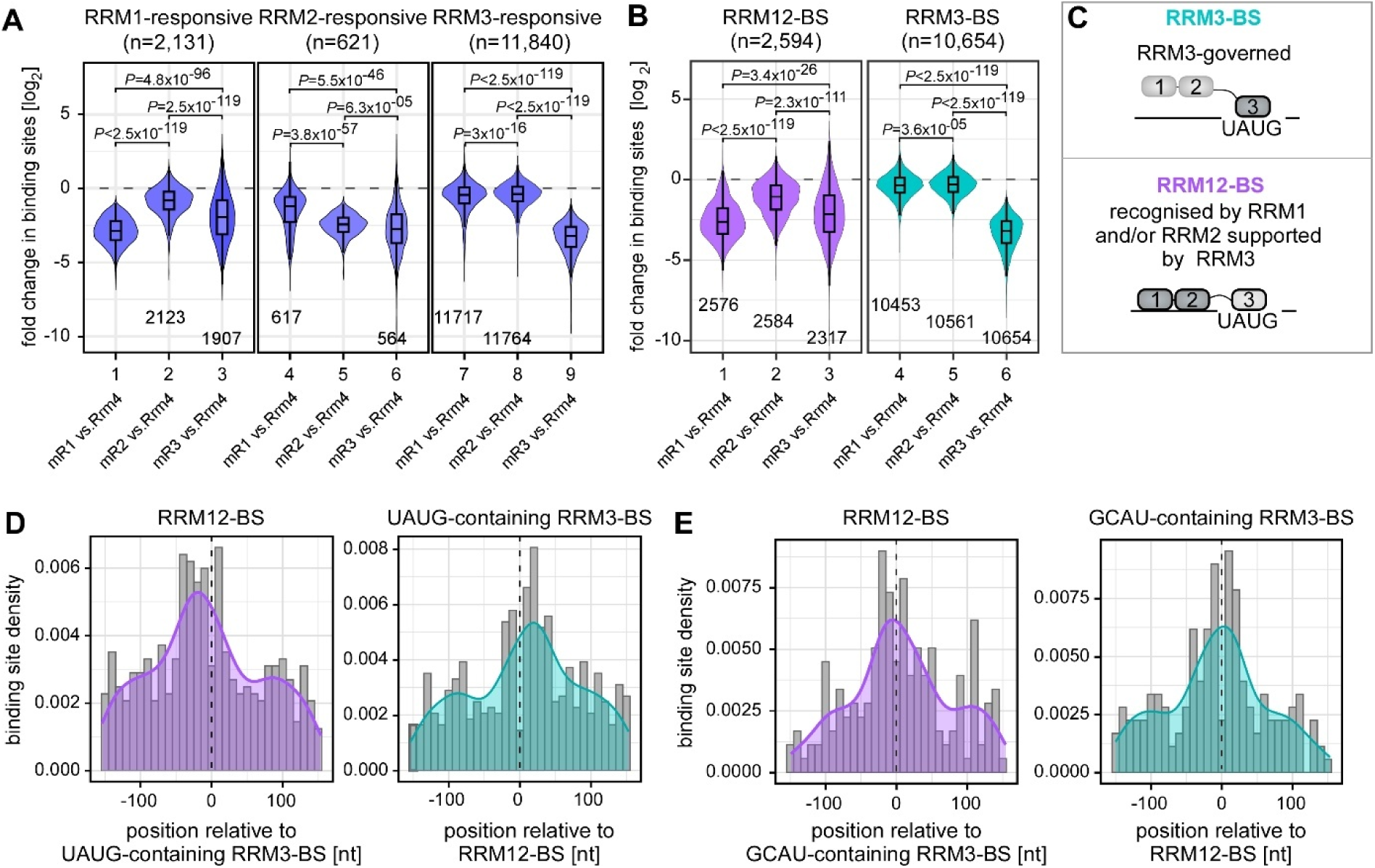
Differential iCLIP2 analysis reveals two types of Rrm4 binding sites. (**A**) Violin plots (numbered at the bottom) showing the quantitative changes in crosslink events at binding sites for RRM1- (left box), RRM2- (center box), and RRM3-responsive sites (right box) in response to mutation in RRM1 (mR1 vs. Rrm4), RRM2 (mR2 vs. Rrm4) and RRM3 (mR3 vs. Rrm4). Boxes represent quartiles, with the center line marking the median. The whiskers extend to the most extreme values within the 1.5x interquartile range. The number of responsive sites measured in the mutant datasets is indicated at the bottom. Statistical significance for changes in responsive sites among the mutant datasets was assessed using the Wilcoxon signed-rank test. (**B**) Violin plots illustrate the quantitative changes in RRM12-BS (purple; left) or RRM3-BS (petrol; right) in mutant datasets. Statistical significance was determined using the Wilcoxon signed-rank test. (**C**) Graphical representation of Rrm4 binding sites. RRM3-BS: Binding sites governed by the third RRM domain, with minimal influence from the RRM1 and RRM2 domains. RRM12-BS: Binding sites recognized by RRM1 and/or RRM2 with often additional support from RRM3. (**D**) Histogram with density curve illustrates the positional distribution of RRM12-BS (left panel) within a 301 nt window centered on RRM3-BS with UAUG motifs or *vice versa* (right panel). (**E**) Same as (D) but using RRM3- BS with GCAU control motifs.

Based on these results, we defined two distinct sets of binding sites for Rrm4: RRM12-BS and RRM3-BS (Fig. 3B-C, Fig. EV3A). In the case of RRM3-BS, binding is mainly governed by the third RRM domain, with the other two contributing only marginally (Fig. 3A-C; Fig. EV3A; n=10,654). In contrast, for RRM12-BS, binding strongly relies on RRM1 and/or RRM2 of the tandem RRM domains, with RRM3 often supporting the interaction (Fig. 3A-C, n=2,594). Mutations in RRM1 show a stronger effect on RRM12-BS than RRM2, underscoring the critical role of RRM1 in recognizing these targets (Fig. 3B).

To test whether the RRMs bind in a defined order on the target mRNAs, we analyzed the positioning of RRM12-BS and RRM3-BS relative to each other in a 301 nt window. Of note, 48% of the RRM12-BS contained at least one RRM3-BS within this window, suggesting that the RRMs bind in the vicinity of each other. Moreover, RRM12-BS tended to accumulate upstream of UAUG-containing RRM3-BS (Fig. 3D-E). This pattern suggests that the RRMs of Rrm4 are positioned in a defined arrangement, with the tandem RRM1 and 2 placed upstream of RRM3 (Fig. 3C). Consistently, the RRM12-BS were enriched predominantly in coding sequences (CDS), whereas RRM3-BS were increasingly found in 3’ UTRs (Fig. EV3B).

In essence, our transcriptome-wide comparative binding analysis highlights the coordination between the RRMs in facilitating Rrm4 binding. Specifically, RRM3 plays a key role in recognizing RRM3-BS, while RRM1 and RRM2 bind together at composite RRM12- BS, which are further supported by RRM3.

### The composite RRM12-BS function as regulatory RNA elements

Consistent with the notion that RRM3 supports the binding of RRM1 and RRM2 at RRM12- BS, most mRNAs with RRM12-BS also harbored at least one RRM3-BS, whereas the latter also frequently occurred alone (Fig. EV4A). Based on this, we categorized target mRNAs as either RRM12-BS targets (n=1,630) or RRM3-BS targets, whereby the latter contained only RRM3-BS but no RRM12-BS (n=2,211; Fig. EV4A). To analyse the regulatory impact of Rrm4 binding to these targets, we performed RNA sequencing (RNA-seq) to measure mRNA levels in the Rrm4-G strain and the corresponding mutant strains. A quantitative comparison revealed that *rrm4Δ*, mR1, mR2, and mR123 mutations resulted in significant changes in mRNA levels, while the mR3 mutation showed only marginal effects (Fig. EV4B). Of note, we observed a clear correlation between targets carrying RRM12-BS and a reduction in relative transcript levels across all mutants with impaired tandem RRM binding (Fig. 4A; Fig. EV4C). Strikingly, this decrease was proportional to the number of RRM12-BS present in target transcripts, indicating an additive effect. Moreover, this pattern was absent in mR3 (Fig. 4A). In contrast, the RRM3-BS targets exhibited no significant changes or additive effects following *rrm4* deletion or any RRM domain mutation (Fig. 4A). This demonstrates that RRM12-BS are functional regulatory RNA elements that link RNA transport and RNA abundance, further underscoring the importance of RRM1 and RRM2. Importantly, despite their functional relevance, the RRM12-BS are less prominent than the RRM3-BS (Fig. EV4F, EV5A-B) and hence easily overlooked in the wild-type Rrm4 iCLIP2 data, underlining the advantage of the comparative iCLIP2 approach in discriminating the effects of the individual RRM.

**Figure 4.**
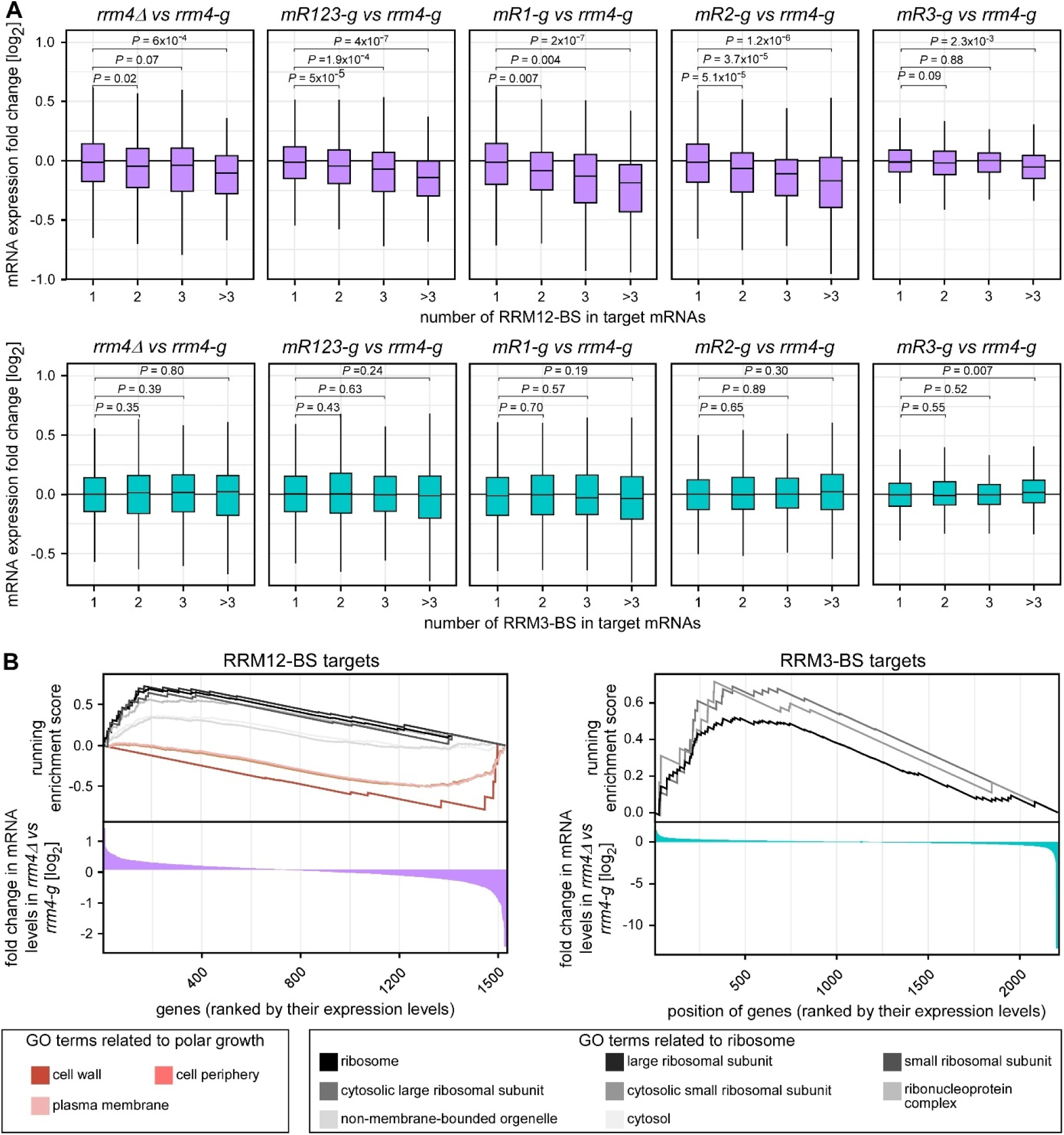
RRM12-BS constitutes functional RNA regulatory elements. (**A**) Box plots illustrate the fold changes in mRNA levels of RRM12-BS targets (purple; top) and RRM3-BS targets (petrol; bottom) in strains lacking the *rrm4* gene (*rrm4Δ*), or those carrying mutations in RRM domains (*mR1-g*, *mR2-g*, *mR3-g, and mR123-g*) compared to wild-type *rrm4* gene (*rrm4-g*). The target mRNAs are grouped based on the number of RRM12-BS (top) or RRM3-BS in them. Statistical significance between groups was assessed using the Wilcoxon signed-rank test. (**B**) Gene set enrichment analysis showing the enrichment of GO terms related to cellular compartments in RRM12-BS targets (left) and RRM3-BS targets (right). Terms associated with polar growth are colored in red shadings, while those related to ribosomes are colored in shades of black. The colored bars below represent the changes in mRNA levels of RRM12-BS targets (purple) and RRM3-BS targets (petrol), ranked in decreasing order by their fold change.

Following up on the functional relevance of tandem RRM binding, an analysis of Gene Ontology (GO) term enrichment revealed that terms related to polar growth, such as cell periphery and plasma membrane, were significantly overrepresented in RRM12-BS targets, whereas no significant enrichment was observed in RRM3-BS targets (Fig. EV4D-E). A clear functional distinction between RRM12-BS and RRM3-BS targets was further supported by a gene set enrichment analysis, which incorporated mRNA level changes following *rrm4* deletion into gene sets based on cellular compartments (Fig. 4B). Importantly, RRM12-BS targets, which exhibited reduced levels in the respective mutants, were strongly enriched for compartments related to cell wall synthesis, cell periphery, and plasma membrane. This was not observed in the case of RRM3-BS targets. Conversely, mRNAs encoding ribosomal proteins were enriched in both RRM12-BS and RRM3-BS targets and showed increased levels in the absence of Rrm4, suggesting a potential compensatory mechanism due to disturbed RNA transport (Fig. 4B). In summary, RRM12-BS exhibit characteristic features of regulatory RNA elements important for Rrm4-dependent transport of cognate target mRNAs during polar growth.

### RRM12-BS targets encode polarity factors, cytoskeletal proteins, and cell wall enzymes

Rrm4 has a well-established role during unipolar hyphal growth. To elucidate the contribution of RRM12-BS to cell polarity – as suggested by the GO analysis – we examined the 1,630 RRM12-BS target mRNAs in greater detail. Generally, hyphal growth is initiated by the subcellular accumulation of polarity factors at the growth pole. These regulators reorganize the cytoskeleton to orchestrate efficient expansion of plasma membrane and cell wall at defined sites (64). Interestingly, within the relevant GO terms *cell periphery*, site of polar growth, and growing tip, we observed several key polarity factors such as small GTPases Cdc42, Rac1, and Rho1/2/4 (Fig. 5A-B, Supplementary Table S7). Consistently, mRNAs encoding cytoskeletal proteins such as F-actin and septin proteins (Cdc3, Cdc11, Cdc12) were RRM12-BS targets (Fig. 5B, Fig. EV5A, Supplementary Table S7). Septin-encoding mRNAs are established Rrm4-dependent cargos, and their endosome-coupled translation is crucial for both the assembly of heterotetrameric complexes and the formation of a septin gradient at the growing pole (36,37).

**Figure 5.**
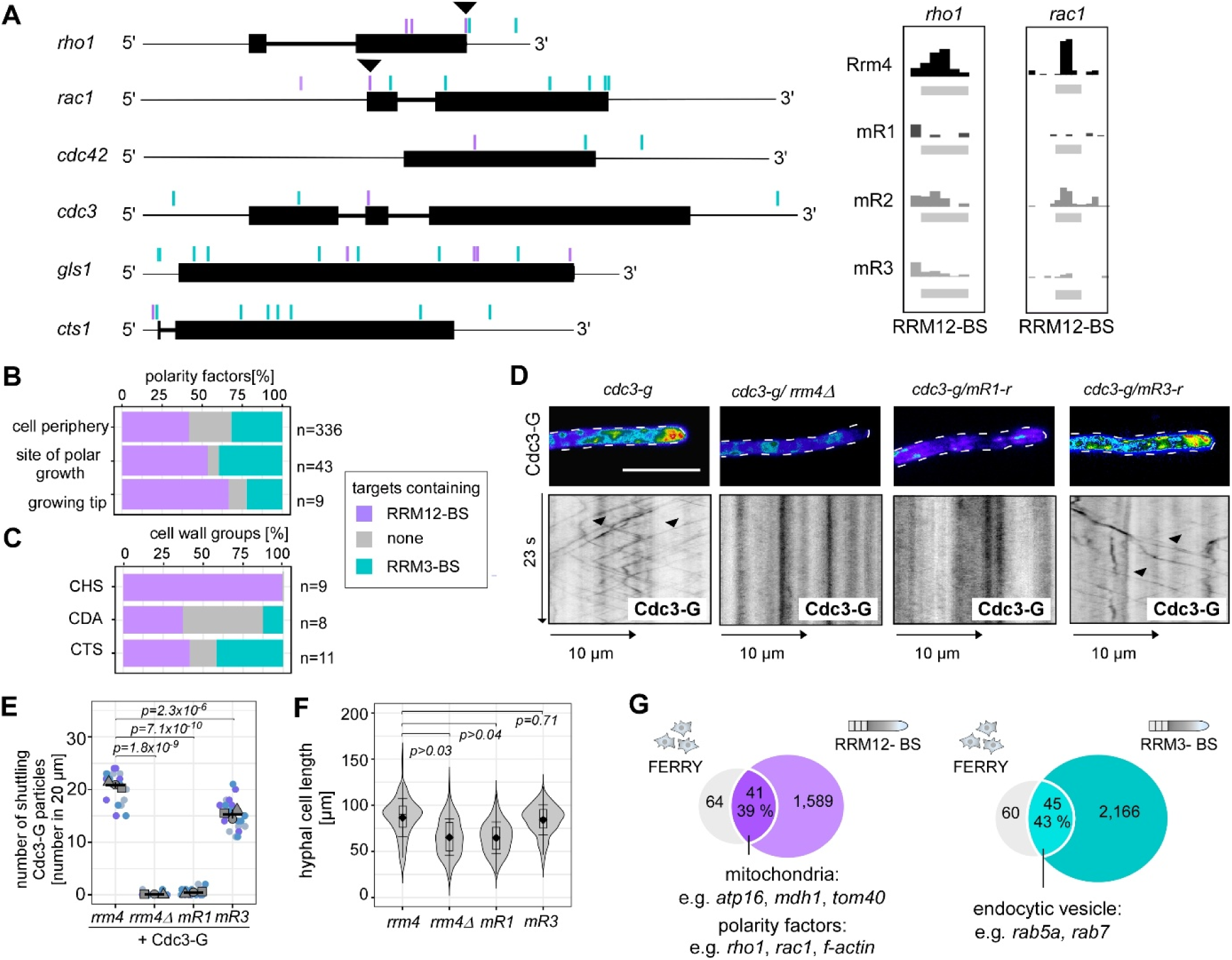
RRM12-BS targets link Rrm4 function to cell wall synthesis, mitochondrial metabolism and polarity. **(A**) Genome browser visualization of RRM12-BS containing target mRNAs, displaying differential Rrm4 binding site profiles. RRM3-BS (petrol) and RRM12-BS (purple) are highlighted, indicating distinct regulatory contributions of individual RRMs. A zoom-in view of selected RRM12-BS regions (arrowheads) is shown on the right, illustrating the crosslink event landscape for Rrm4, mR1, mR2, and mR3 (group autoscale). (**B**) Bar diagram showing the percentage of target mRNAs containing RRM12-BS, RRM3-BS or none within GO terms associated with polar growth and polarity factors: *cell periphery* (GO:0071944, n=336), *site of polar growth* (GO:0030427, n=43), and *growing cell tip* (GO:0035838, n=9). The entries (n) of the GO terms are indicated. **(C**) Bar diagram as in (B) exhibiting cell wall remodeling enzymes categorized in chitin synthase (CHS, n=9), chitin deacetylase (CDA, n=8), and degradation (e.g. chitinases, CTS, n=11). (**D**) The upper panel represents false color imaging of Cdc3-G fluorescence of the growing cell tip (6. h. i., size bar: 10 µm) in diverse Rrm4 background strains. The lower panel shows the respective kymographs of shuttling Cdc3-G particles. (**E**) Quantification of Cdc3-G shuttling particles (Student’s *t*-test, unpaired; based on the means of the three biological replicates, replicate 1=circle; replicate 2=triangle, replicate 3=square; N > 20). (**F**) Violin plot of hyphal cell length (6. h. p. i., Student’s *t*-test, unpaired; three biological replicates, N > 200). (**G**) Venn diagrams of potential FERRY targets and RRM12-BS (purple) and RRM3-BS (petrol) containing targets.

In line with polar cell wall expansion as a critical process in hyphal growth, RRM12-BS targets were strongly enriched for mRNAs encoding enzymes for cell wall synthesis and remodeling. This included all chitin synthases, Chs1-7 and Mcs1, which are involved in the synthesis of cell wall chitin (Fig. 5C, Fig. EV5B, Supplementary Table S7). Furthermore, transcripts coding for enzymes acting in cell wall remodeling (Fig. 5A-C) included 1,3-β-glucan synthase Gls1 (65,66), three chitin deacetylases (Cda1, Cda6-7), chitinases Cts1 and Cts2, and three glycoside hydrolases (Gla5-7) (67) were also RRM12-BS targets. Noteworthy, we could previously show that *cts1* mRNA is a known Rrm4 target and the unconventional secretion of Cts1 is Rrm4-dependent (68). Together, Rrm4-mediated mRNA transport appears to coordinate cytoskeletal dynamics and cell wall remodeling required for polar growth preferentially via the coordinated action of the tandem domains RRM1 and 2 at RRM12-BS.

For experimental verification of these observations, we chose septin Cdc3 (Fig. 5A). The corresponding mRNA contains one RRM12-BS and three RRM3-BS, providing an ideal target to investigate how variations in RRM-dependent interactions affect the subcellular localization of functional Cdc3 protein (Fig. 5D-E). Consistent with previous reports, Cdc3 fused to Gfp (Cdc3-G, Supplementary Table S1) was barely detectable on endosomes in *rrm4Δ* and RRM1 mutants (mR1), and the gradient at the tip was lost (Fig. 5D-E) (32,36,37). Mutations in RRM3 (mR3) caused a slight but significant reduction in endosomal shuttling and gradient formation (Fig. 5D-E), supporting the idea that RRM3 enhances RRM1 binding, while the tandem RRMs are essential for RNA transport. Accordingly, cell length that depends on cell wall synthesis was reduced in strains expressing mR1, but not mR3 (Fig. 5F). In essence, RRM12-BS targets encode factors contributing to polar growth at all levels, from initial regulation to cell wall remodeling, including a novel extensive link to cell wall synthesis.

### A core set of endosomal transport target mRNAs is evolutionarily conserved from fungi to humans

We leveraged evidence from mammalian cells to investigate the evolutionary conservation of endosomal mRNA transport and hence its relevance beyond *U. maydis*. In neurons, the FERRY complex links mRNAs to early endosomes and using it as a bait in pull-down experiments, 252 target mRNAs were identified in human HEK293 cells (22). Reciprocal BLAST analysis identified 122 unambiguous orthologs in *U. maydis*, the majority of which was bound by Rrm4 (n=105, 86%; Fig. EV5C-D, Supplementary Table S5). When dissecting the overlap in Rrm4’s binding sets (RRM12-BS vs RRM3-BS), the shared pool of targets was approximately evenly divided, with 40% bound by RRM12-BS and 40% by RRM3-BS (Fig. EV5D, Supplementary Table S5). However, RRM3-BS targets were enriched for endocytic vesicles and intracellular structures. In contrast, RRM12-BS targets were enriched for the GO terms plasma membrane and mitochondrial proteins such as Atp16 and Mdh1. Interestingly, this data set also contained small GTPases such as Rho1 and Rac1 (Fig. 5G, Supplementary Table S5). In essence, a comparison of mRNA targets of the endosomal transport machinery from fungi and humans identified an evolutionarily conserved subset of targets involved in mitochondrial homeostasis and determination of polarity.

## Discussion

RNA-binding proteins often use multiple RBDs to recognize defined RNA regulons. To understand their precise regulatory capacity, it is essential to dissect how individual RBDs distinguish essential from accessory interactions (Fig. 6A) (8,9,64). Here, we established a multifaceted comparative iCLIP2 approach to decipher the interplay of multiple RRMs present in the endosomal mRNA transporter Rrm4. We succeeded in differentiating functional from accessory binding sites and thereby disclosed new RNA transport regulons. Our unique study combining single-nucleotide UV crosslinking of RBD mutants with transcriptomics and cell biology might serve as a blueprint to tackle the RNA binding code in general and advance *in vivo* RNA biochemistry.

**Figure 6.**
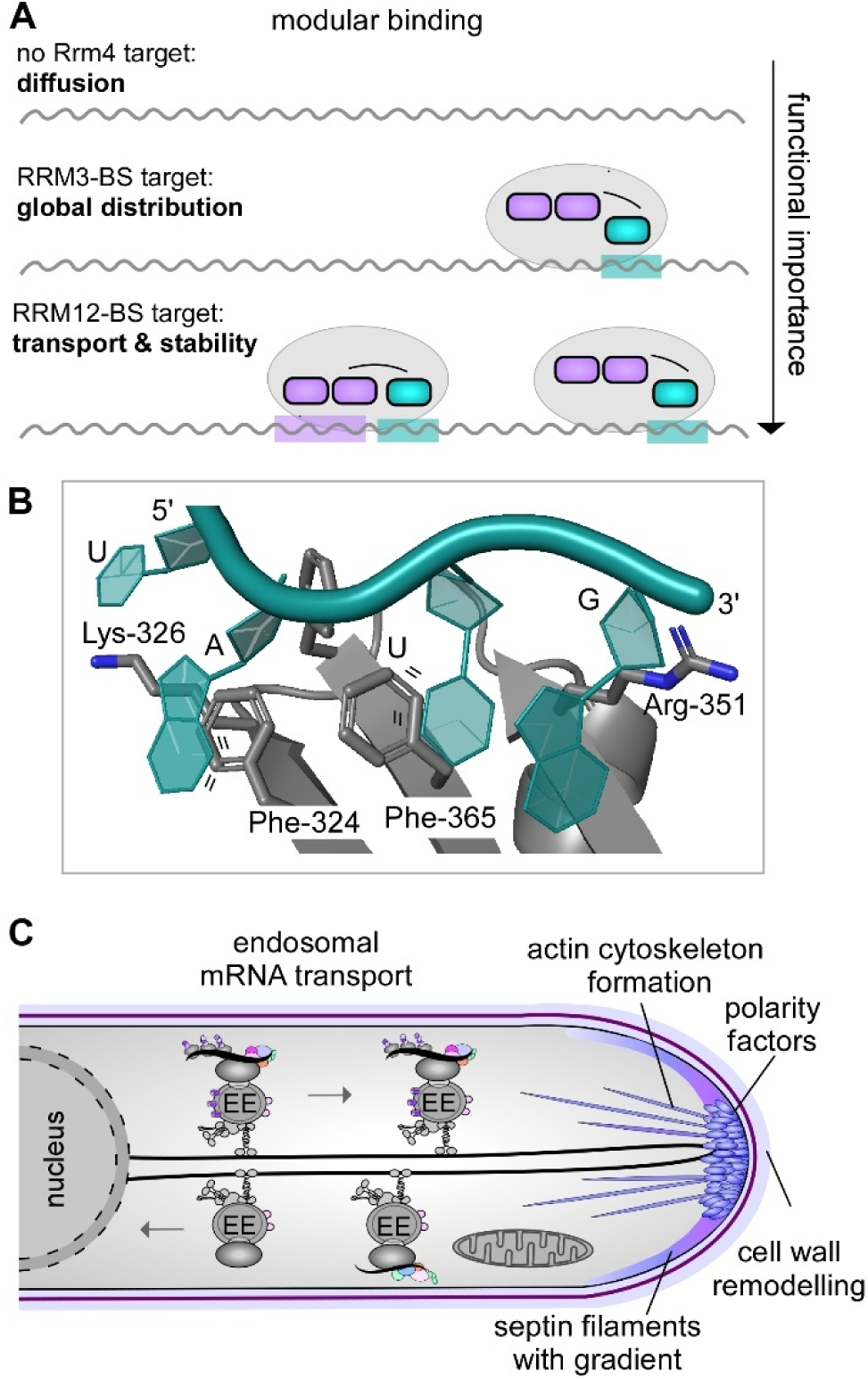
The comparative iCLIP2 approach identifies accessory and key binding sites of the multi-RRM protein Rrm4. (**A**) Modular binding mode of the multi-RRM protein Rrm4. RRM3 (petrol) and its specific binding sites (RRM3-BS, petrol) are essential for the uniform distribution of RRM3-BS-containing targets throughout the hyphae. The tandem RRMs (RRM1 and RRM2=purple) are crucial in regulating the transport and stability of RRM12-BS-containing mRNAs, which encode proteins that specifically localize to the cell tip, driving cell wall expansion and polarity growth. The regulation of these RRM12-BS targets is primarily controlled by RRM1 and/or RRM2, with additional support from RRM3. Importantly, the abundance of RRM12-BS within a target correlates with a significant effect on its mRNA abundance, highlighting the intricate interplay between these binding modes and cellular architecture. **(B**) AlphaFold3 model of RRM3/UAUG interaction. For prediction, full-length Rrm4 was offered. Among the three RRM domains, RRM3 was selected. Potential π-stacking events are indicated (lines). **(C**) Schematic representation of endosomal mRNA transport in *U. maydis*. Functional regulated RRM12-BS containing regulons such as polarity factors, septin-gradient and cytoskeleton formation, cell wall remodeling, as well as mitochondria, are highlighted

### RRMs recognize defined sets of binding sites to distinguish accessory from main functions

Currently, most transcriptome-wide RNA binding studies, which use iCLIP2 and variations, employ wild-type RBPs and provide a wealth of information concerning binding motifs, binding sites, and target mRNAs (7,8). For instance, PAR-CLIP/CRAC approaches of the multi-RRM RBPs, HuR and HuD, revealed AU-rich binding motifs within hundreds of target mRNAs (17,69). Similarly, our previous iCLIP study of wild-type Rrm4 identified *bona fide* mRNA targets and revealed UAUG as the dominant binding motif (34). However, the ultimate challenge is to unravel how individual RBDs drive the specific recognition of functionally relevant regulatory RNA elements (Fig. 6A) (10,11,70).

An attractive approach is the comparison of wild-type and RBD mutants. For example, iCLIP2 was used to compare the RNA-binding ubiquitin E3 ligase TRIM25 with its RBD mutants, unraveling their importance in RNA binding (71). Here, we advanced this concept further. By analyzing RBD mutant variants of the multi-RRM protein Rrm4, we determined their distinct binding sites and functions with high resolution. We refined our iCLIP2 protocol to achieve high-quality data (40), enabling precise detection of even subtle changes in RBD mutants with up to 60% reduced binding.

Advanced differential binding analysis (51) revealed two distinct Rrm4 binding sets validated by target transcript expression profiling: functionally critical sites primarily recognized by the tandem RRMs RRM1 and/or RRM2 (RRM12-BS, Fig. 6A) differ from accessory sites predominantly bound by RRM3 (RRM3-BS, Fig. 6A). Intriguingly, RRM3 supported the binding activity of RRM1 and RRM2, emphasizing its auxiliary role in overall RNA binding (Fig. 6A). Mutations in RRM3 caused a subtle yet significant shift in bulk mRNA distribution as well as Cdc3 shuttling and localization but did not disrupt unipolar growth. This indicates that while RRM3 is supportive, RRM1 and RRM2 mediate the functionally critical RNA contacts (Fig. 6A). These findings align with *in vitro* studies of the multi-RRM RNA transporter HuD, where RRM1 and RRM2 mediate RNA binding, and RRM3 enhances binding affinity (21,72).

While solving the multi-RRM code, we discovered that the binding of RRM3 to UAUG resulted in the most prominent UV crosslinks, despite their limited functional significance as seen for septin Cdc3. A plausible explanation is a quantitative UV crosslinking bias due to π- stacking interactions between aromatic amino acids and predominantly U or G nucleotides (73). RRM3 contains four aromatic amino acids (Fig. EV1A) known for efficient crosslinking to uracil (74). Consistently, an AlphaFold3 prediction indicates potential π-stacking interactions between the phenylalanine side chains of RRM3 and both uracils of UAUG (Fig. 6B) (3). This nicely illustrates that pronounced UV crosslink events may not reflect the most functionally significant binding sites *in vivo*.

### RRM12-BS exhibit characteristic features of regulatory RNA elements

Our transcriptome-wide binding analysis successfully distinguished functionally relevant Rrm4 binding sites (RRM12-BS) from accessory sites (RRM3-BS), emphasizing the central role of RRM1 and RRM2 in target recognition and functional regulation. Loss-of-function mutants showed a significant reduction in mRNA levels of RRM12-BS targets and an increasing number of RRM12-BS sites amplified this effect, indicating a direct link between binding site abundance and mRNA levels. This highlights RRM12-BS as a hallmark regulatory RNA element. Our results are consistent with current views reporting that vesicle-mediated transport safeguards mRNAs from degradation during transit in mammals (22,24). Such a protective function was also described for the RBP Oskar from *D. melanogaster* that shields *nanos* mRNA during localization in embryos (75). We, therefore, propose that Rrm4-mediated transport protects RRM12-BS-containing mRNAs from cytoplasmic degradation, linking this process to changes in mRNA abundance. This is in accordance with a global link between mRNA stability and defined subcellular localization that was recently disclosed in neurons (29,76,77). Overall, we uncovered distinct functional and spatial specificities of RRM1 and RRM2 compared to RRM3 (Fig. 6A), providing new insights into the nuanced regulation of RNA binding, transport, and stability.

### Target mRNAs encode factors orchestrating polar growth

mRNA translocation by vesicle hitchhiking is a widespread mechanism and mRNAs encoding mitochondrial proteins have already been identified as evolutionarily conserved cargos of this process (Fig. 6C) (3,22,24,26,27,30,34,78). Consistently, RRM12-BS targets encode subunits present in all five respiratory chain complexes and as anticipated, human FERRY and *U. maydis* Rrm4 share mitochondrial targets. Intriguingly, a second common mRNA regulon transported by both endosomal transport machinery are mRNAs encoding small GTPases such as Rac1 and polarity factors (22–24). This might be particularly important to determine fungal and mammalian cell polarity (Fig. 6C) (3,79–81).

Besides the FERRY complex, other RBPs are most likely involved in vesicle-mediated mRNA transport. TDP-43-containing mRNPs, for example, are transported by lysosome hitchhiking in axons (26,78) and TDP-43 also binds mRNAs encoding polarity factors such as *RAC1* mRNA (82–84). Thereby, abberant TDP-43 might contribute to neuronal diseases such as amyotrophic lateral sclerosis and frontotemporal lobar degeneration (85).

Finally, RRM12-BS containing target mRNAs were strongly enriched in cell wall modifying enzymes. These most likely foster local cell wall remodeling (Fig. 6C) (65,66), disclosing a novel link between the coordination of cell wall synthesis and endosomal mRNA transport. Fundamentally, our study not only identifies novel targets involved in cell wall synthesis and modification but also uncovers the core targets of endosomal mRNA transport, evolutionarily conserved across polarized eukaryotic cells.

## Conclusions

Our powerful comparative iCLIP2 approach assigned specific functions to individual RRMs. Thereby, accessory sites were distinguished from functional sites, which in turn orchestrate spatiotemporal expression of distinct mRNA regulons. Our findings underscore the complexity of RNA recognition, providing a new perspective beyond simple target identification toward understanding global RNA binding dynamics. Decoding modular RRM binding by iCLIP2 revealed a novel connection between endosomal transport and the subcellular localization of polarity factors – a pathway seemingly conserved in highly polarized cells such as fungal hyphae and neurons. This insight broadens our understanding of membrane-coupled mRNA transport, with broad implications from fighting fungal pathogens to the development of novel therapeutic approaches in neuronal diseases (5).

## Data availability

RNA-seq and iCLIP2 data are available in the NCBI Gene Expression Omnibus (GEO). The iCLIP2 data for wild-type Rrm4 can be accessed under GSE273496, while the mutant Rrm4 variants are available under GSE273495 and require the security token xx for access. The RNA-seq data are available under GSE288123, with anonymous reviewer access provided via the security token xx.

## Acknowledgments

We sincerely thank Drs. Heiner Schaal, Florian Altegoer, Kerstin Schipper, and lab members for their critical reading of the manuscript and valuable feedback. We are particularly grateful to Dr. Florian Altegoer for his expert guidance on structural prediction and to Dr. Mirko Brüggemann and members of the Zarnack group for insightful discussions on binding site definition and differential binding analysis. We appreciate the dedicated support of Maximilian Neumann, Marie Wittmann, and Caroline Downes in plasmid generation and Ute Gengenbacher and Simone Esch for their exceptional technical assistance regarding strain generation. We extend our thanks to Dr. Maria Méndez-Lago and the IMB Genomics Core Facility in Mainz for their outstanding support with iCLIP2 and mRNA library sequencing, as well as for granting access to their Illumina NextSeq 500 platform.

## Author contributions

NKS, SS, KZ, JK, and MF designed this study and analyzed the data. NKS performed wet-lab experiments supported by JK and KM. *U*. *maydis* strains were planned by NKS, KM, and MF. SS, AB, NKS, and KZ performed the bioinformatic analysis. NKS, SS, KZ, and MF drafted and revised the manuscript with input from all co-authors. MF, KZ, and JK contributed funding and resources.

## Expanded View Figures

**Figure EV1.**
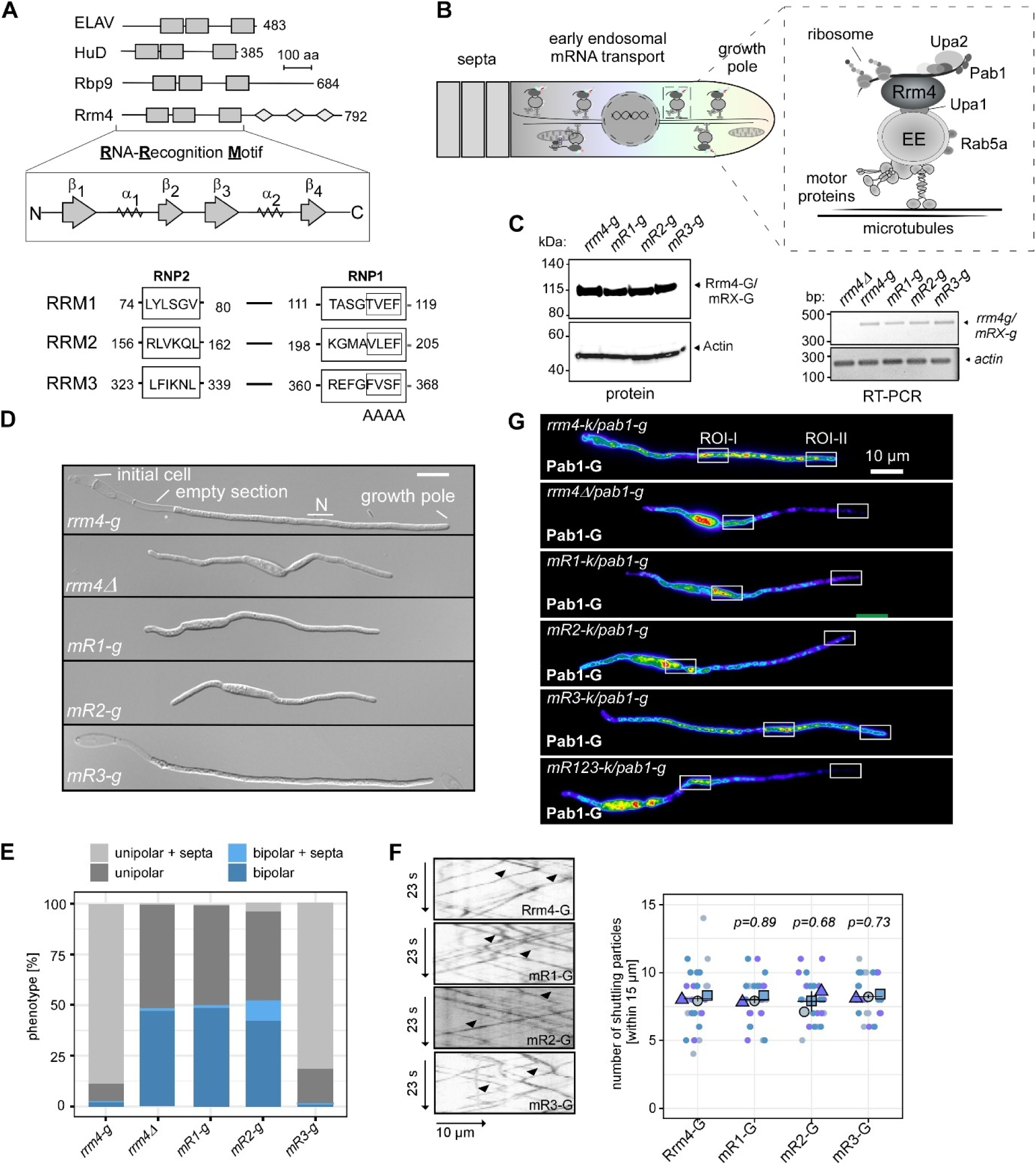
Mutations in RRM1 and RRM2 cause loss-of-function. (**A**) Schematic representation of diverse multi-RRM RNA binding proteins: ELAV (*Drosophila melanogaster*), HuD (*Homo sapiens*), Rbp9 (*Drosophila melanogaster*), and Rrm4 (*Ustilago maydis*) and a detailed view of the structural architecture of a typical *RNA-recognition motif* (RRM). (**B**) Schematic representation of the current endosomal mRNA transport model *in U. maydis.* The key RBP Rrm4 (RNA recognition motif 4, dark grey), accessory RBP Pab1 (Poly(A) binding protein 1, anthracite) and associated ribosomes are indicated. In addition, the mRNP scaffold protein Upa2, the anchor protein Upa1, and the early endosomal marker protein Rab5a are labeled. (**C**) Western blot (left) analysis of Rrm4-G (112 kDa) and the respective RRM mutants (mR1-mR3-G: 112 kDa). Rrm4-G/mRX-G full-length band is indicated by an arrow. Actin protein (45 kDa) serves as a loading control. (right) RT-PCR of *rrm4* mRNA and the respective mutant versions (mR1-mR3). The *actin* mRNA serves as constitutive control for the analysis of equal RNA amounts. (**D**) DIC pictures of hyphae (6 h. p. i, size bar, 10 µm). (**E**) A stacked bar graph of quantified polarity phenotypes of hyphae: unipolarity, bipolarity, and septum insertion were counted (6 h. p. i., three biological replicates, N>100). (**F**) Kymographs (inverted fluorescence images) of hyphae (6 h. p. i.) showing Rrm4-G and respective mutants (mR1-mR3-G). Right side, a scatter plot of the number of shuttling particles measured within 15 µm sections (Student’s *t*-test, unpaired; three biological replicates indicated by different blue shadings, N=30). (**G**) False color imaging of Pab1-G fluorescence in hyphae (6. h. p. i.). Regions of interest (ROI) are marked by a rectangle (size bar, 10 µm).

**Figure EV2.**
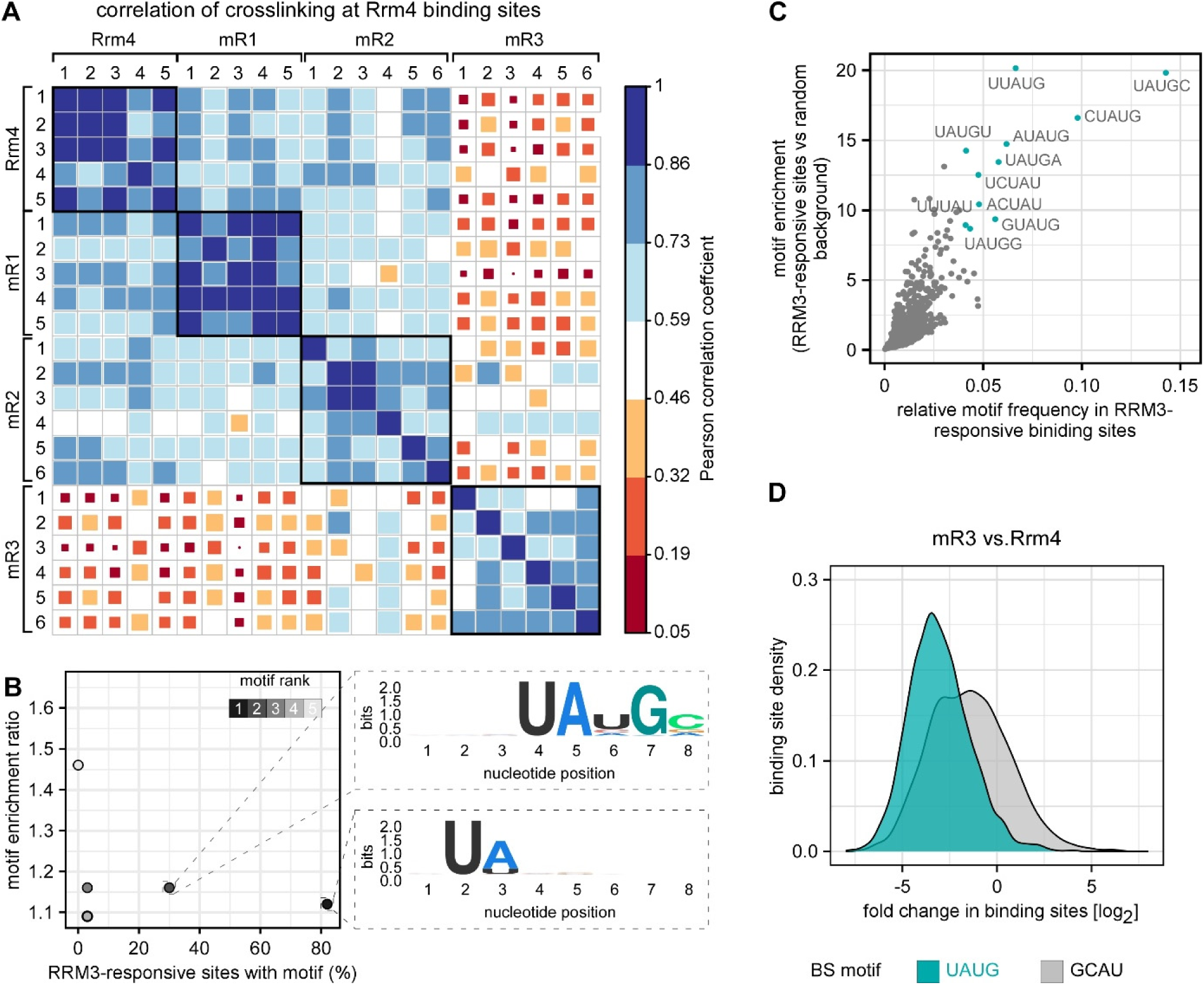
UAUG is a prominent RRM3 binding motif. (**A**) Correlation of crosslinking events per Rrm4 binding sites across iCLIP2 (Rrm4; mR1; mR2; mR3) and their respective replicates (minimum of five replicates). The size and color intensity of the square represents the absolute value of the Pearson correlation coefficient. (**B**) *De novo* motif discovery analysis on RRM3-responsive binding sites. Scatter plot compares the relative enrichment ratio of each motif with its percentage in the RRM3-responsive binding sites. The enlarged section displays the sequence logos of the top two ranked motifs. The motifs were colored according to their ranks. (**C**) Motif enrichment in the RRM3-responsive sites. Pentamer frequencies were calculated by comparing the RRM3-responsive motifs to a random genome background. Highly enriched motifs were highlighted in petrol. (**D**) Density plot showing the distribution of binding site fold change [log_2_] for UAUG-containing sites (petrol) and GCAU-containing sites (control motif; grey) in the mR1 vs Rrm4 dataset.

**Figure EV3.**
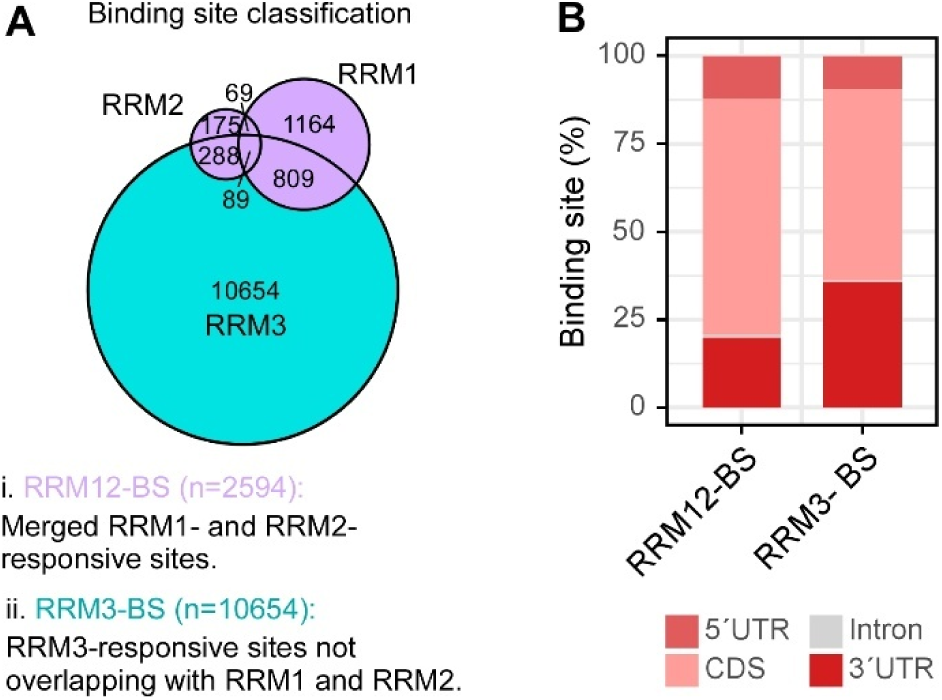
RRM12-BS differ from RRM3-BS. (**A**) Overview of Rrm4 binding site classification. RRM1- and RRM2-responsive sites were combined and categorized as RRM12-BS (purple region in the Venn diagram). RRM3- responsive sites that do not overlap with RRM1- and RRM2-responsive sites were classified as RRM3-BS (petrol colored region in the Venn diagram). (**B**) Distribution of RRM12-BS and RRM3-BS across transcript regions. (CDS- coding sequence; UTR-untranslated region).

**Figure EV4.**
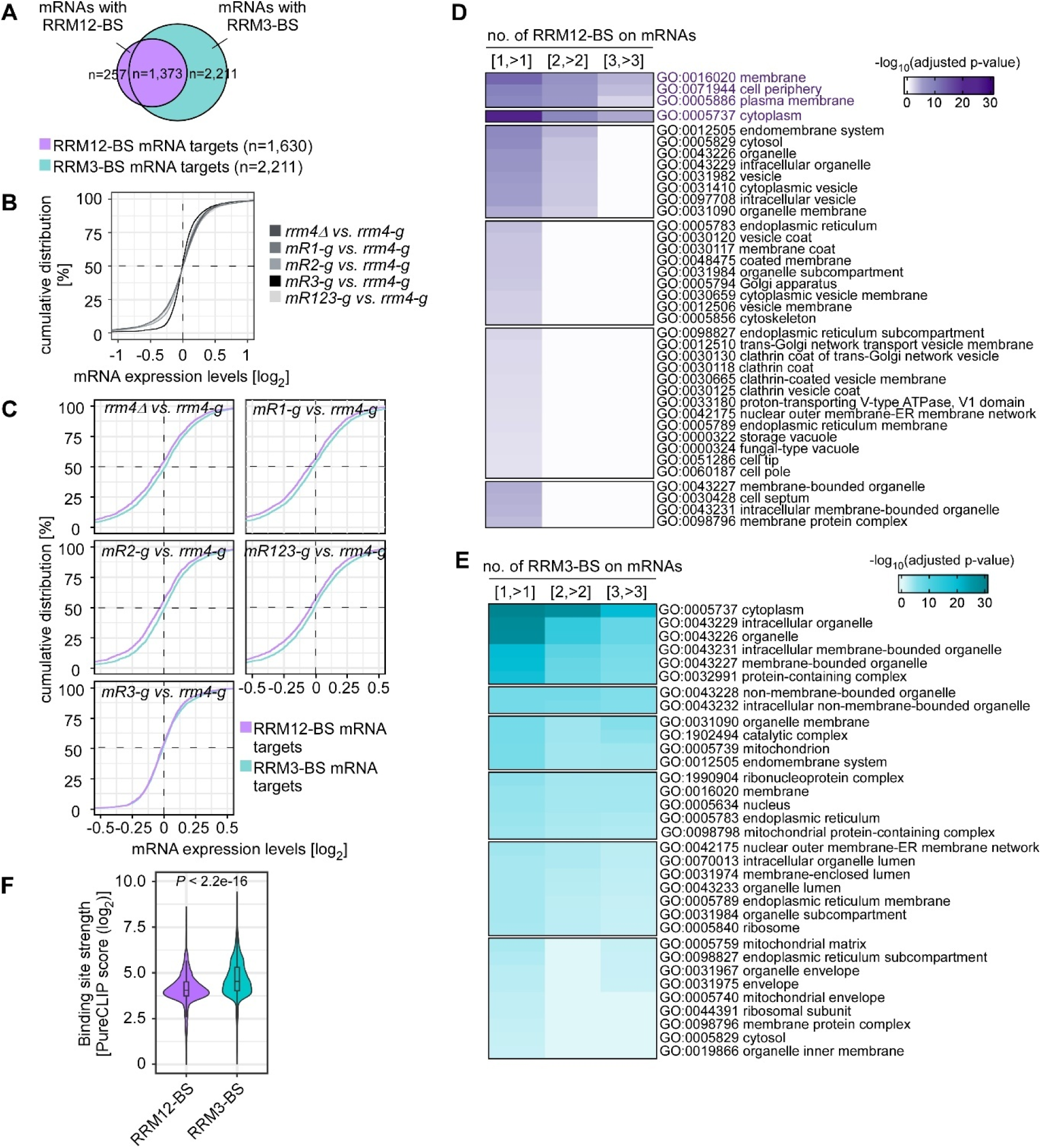
RRM12-BS target exhibit reduced mRNA amounts if Rrm4 function is affected. (**A**) Venn diagram comparing the overlap between RRM12-BS and RRM3-BS mRNA. mRNA targets were assigned hierarchically. (**B**) Cumulative fraction of all hyphae expressed transcripts (n=6278) with the given difference in mRNA expression levels [log_2_] for *rrm4Δ vs. rrm4-g*, *mR1-g vs. rrm4-g*, *mR2-g vs. rrm4-g*, *mR3-g vs. rrm4-g* and *mR123-g vs. rrm4-g* datasets. **(C)** Similar to B, for RRM12-BS mRNA targets (n=1630; purple) and RRM3-BS mRNA targets (n=2211; petrol). (**D-E**) Heatmap of gene ontology term (GO term) analysis (57) for RRM12-BS targets (D) and RRM3-BS targets (E). The color scale represents the -log_10_ (adjusted p-values). Categories indicate mRNA targets with at least one binding site [1, >1], at least two binding sites [2,>2], or at least three binding sites [3,>3]. (**F**) Violin plot illustrates the binding site strength (log_2_-transformed PureCLIP score) of RRM12-BS (purple) or RRM3- BS (petrol) in the Rrm4-iCLIP2 dataset. Statistical significance was calculated using the Wilcoxon signed-rank test.

**Figure EV5.**
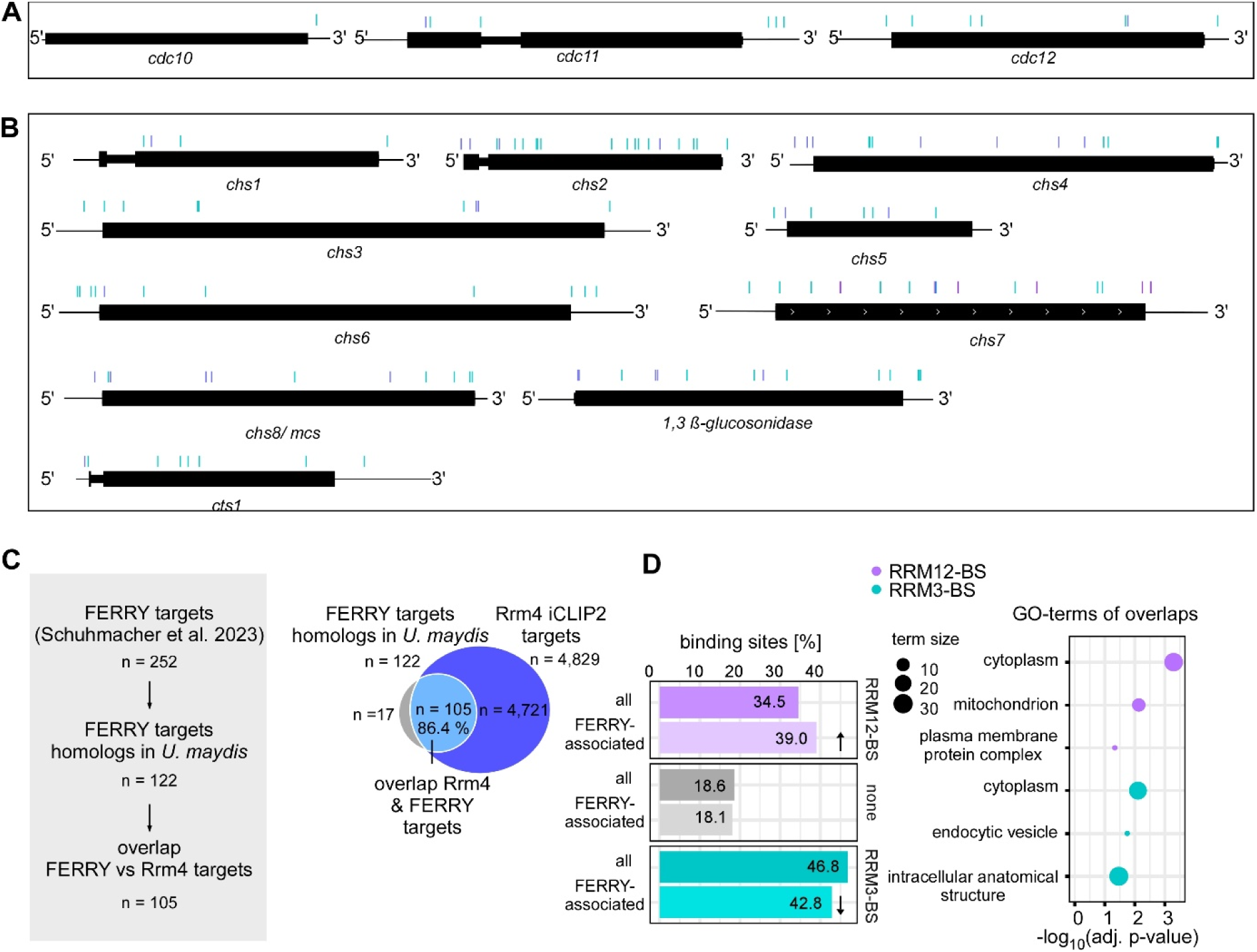
Endosomal mRNA targets are conserved in fungi and mammals. (**A**) Differential binding site profile including RRM3-BS (petrol) and RRM12-BS (purple) of septin mRNAs (*cdc10*, *cdc11*, *cdc12,* Supplementary Table S7-8). (**B**) Genome browser viewer (as in A) of mRNAs encoding enzymes involved in cell wall synthesis (CHS group see Fig. 5, Supplementary Table S7-8). (**C**) Schematical workflow representation and respective overlap between FERRY-associated mRNA (22) and Rrm4-bound targets (light blue; n=105, Supplementary Table S5). (**D**) Percentage of distribution of RRM12-BS-, RRM3-BS-, and none BS- containing targets in overlapping category (FERRY-associated) compared to overall Rrm4- binding targets (all). Right side GO term analysis of overlapping targets: RRM12-BS and FERRY (purple, Supplementary Table S5); RRM3-BS and FERRY (petrol, Supplementary Table S5) performed by gProfiler analysis (57).

