## Supplementary Table S1-4 for "Dissecting the RNA binding capacity of the multi-RRM protein Rrm4 essential for endosomal mRNA transport"

**Addition files:**

Supplementary Table S5 FERRY/ Rrm4 comparison given as a separate Microsoft Excel file

Supplementary Table S6 RRM-responsive side given as a separate Microsoft Excel file

Supplementary Table S7 RRM12-BS given as a separate Microsoft Excel file

Supplementary Table S8 RRM3-BS given as a separate Microsoft Excel file

### Table S1. Description of *U. maydis* strains used in this study

| Strain name with code | Locus | Progenitor strain | Short description |
|---|---|---|---|
| AB33 (UMa133) | *b* | FB2 | *Promotor-nar:bW2bE1*, expression of active b heterodimer under control of the *nar1* promoter, strain grows filamentous upon nitrogen change (Brachmann et al., 2001) |
| AB33_rrm4Δ (Uma273) | *rrm4* | AB33 | carrying a deletion of *rrm4* (Becht et al., 2005) |
| AB33_rrm4-egfp (UMa274) | *rrm4* | AB33rrm4Δ | expressing Rrm4 C-terminally fused to eGfp, endogenous promotor (Becht et al., 2005) |
| AB33_mR1-egfp (UMa311) | *rrm4* | AB33rrm4Δ | expressing a mutated version Rrm4 C-terminally fused to eGfp mutation in RRM1: TVEF (116-119 aa) to AAAA, endogenous promotor (Becht et al., 2006) |
| AB33_mR2-egfp (UMa312) | *rrm4* | AB33rrm4Δ | expressing a mutated version Rrm4 C-terminally fused to eGfp mutation in RRM2: VLEF (201-205 aa) to AAAA, endogenous promotor (this study, plasmid: pUMa636) |
| AB33_mR3-egfp (UMa313) | *rrm4* | AB33rrm4Δ | expressing a mutated version Rrm4 C-terminally fused to eGfp mutation in RRM3: FVSF (362-368 aa) to AAAA, endogenous promotor (Becht et al., 2006) |
| AB33_mR123-egfp (UMa276) | *rrm4* | AB33rrm4Δ | Expressing a triple mutation Rrm4 C-terminally fused to eGfp mutation in RRM1: TVEF (116-119 aa) to AAA; in RRM2: VLEF (201-205 aa) to AAAA, in RRM3: FVSF (362-368 aa) to AAAA, endogenous promotor (Olgeiser et al., 2019) |
| AB33_rrm4-mKate2/pab1-gfp (UMa3334) | *rrm4* *pab1* | AB33 Pab1-egfp-nat/ rrm4Δ (UMa472) | expressing Pab1 C-terminally fused to eGfp and a mutated version of Rrm4 C-terminally fused to mKate2, endogenous promotor (this study: plasmid: pUMa4698) |
| AB33_mR1-mKate2/pab1-gfp (UMa3268) | *rrm4* *pab1* | AB33 Pab1-egfp-nat/ rrm4Δ (UMa472) | expressing Pab1 C-terminally fused to eGfp and a mutated version of Rrm4 C-terminally fused to mKate2; mutation in RRM1: TVEF (116-119 aa) to AAA, endogenous promotor (this study, plasmid: pUMa4569) |
| AB33_mR2-mKate2/pab1-gfp (UL172) | *rrm4* *pab1* | AB33 Pab1-egfp-nat/ rrm4Δ (UMa472) | expressing Pab1 C-terminally fused to eGfp and a mutated version of Rrm4 C-terminally fused to mKate2; mutation in RRM2: VLEF (201-205 aa) to AAAA, endogenous promotor (this study, plasmid: pUL_0192) |
| AB33_mR3-mKate2/pab1-gfp (UL173) | *rrm4* *pab1* | AB33 Pab1-egfp-nat/ rrm4Δ (UMa472) | expressing Pab1 C-terminally fused to eGfp and a mutated version of Rrm4 C-terminally fused to mKate2; mutation in RRM3: FVSF (362-368 aa) to AAAA, endogenous promotor (this study, plasmid: pUL_0193) |
| AB33_mR123-mKate2/pab1-gfp (UMa3270) | *rrm4* *pab1* | AB33 Pab1-egfp-nat/ rrm4Δ (UMa472) | expressing Pab1 C-terminally fused to eGfp and a mutanted version of Rrm4 C-terminally fused to mKate2; mutation in RRM1: TVEF (116-119 aa) to AAA; in RRM2: VLEF (201-205 aa) to AAAA, in RRM3: FVSF (362-368 aa) to AAAA, endogenous promotor (this study plasmid: pUMa4571) |
| AB33_gfp-cdc3 (UMa449) | *cdc3* | AB33 | expressing Cdc3 N-terminally fused to eGfp, endogenous promotor (Baumann et al., 2014) |

| | | | |
|---|---|---|---|
| AB33_gfp-cdc3/rrm4Δ (UMa462) | *rrm4* *cdc3* | AB33_gfp-cdc3 | expressing Cdc3 N-terminally fused to eGfp in an rrm4Δ background, endogenous promotor (Baumann et al., 2014) |
| AB33_Gfp-cdc3-/mR1-rfp- (UMa836) | *rrm4* *cdc3* | AB33_gfp-cdc3 | expressing Cdc3 N-terminally fused to eGfp and a mutated version of Rrm4 C-terminally fused to Rfp; mutation in RRM1: TVEF (116-119 aa) to AAA, endogenous promotor (Baumann et al., 2014) |
| AB33_Gfp-cdc3/mR3-rfp (UMa940) | *rrm4* *cdc3* | AB33_gfp-cdc3 | expressing Cdc3 N-terminally fused to eGfp and a mutated version of Rrm4 C-terminally fused to Rfp; mutation in RRM3: FVSF (362-368 aa) to AAAA, endogenous promotor (this study plasmid: pUMa1703) |

### Table S2. Description of plasmids generated for *U. maydis* strain generation

| Plasmid | Plasmid ID | Resistance cassette | Short description |
|---|---|---|---|
| Prrm4:Rrm4_mR2 _eGfp_ Nat | pUMa636 | NatR (nourseothricin resistance) | Plasmid vector for generating a mutant version of *rrm4* C-terminal fused with eGfp. Mutation in RRM2 -RNP1: VLEF (201-205 aa) to AAAA, construct under control of the endogenous promotor of *rrm4*. |
| Prrm4:Rrm4_mR2_ mKate2_HA_ G418R | pUL192 | genitR (G418 resistance) | Plasmid vector for generating a mutant version of *rrm4* C-terminal fused with mKate2_HA. Mutation in RRM2 -RNP1: VLEF (201-205 aa) to AAAA, construct under control of the endogenous promotor of *rrm4*. |
| Prrm4:Rrm4_mR3_ mKate2_HA_ G418R | pUL193 | genitR (G418 resistance) | Plasmid vector for generating a mutant version of *rrm4* C-terminal fused with mKate2_HA. Mutation in RRM3: FVSF (362-368 aa) to AAAA, construct under control of the endogenous promotor of *rrm4*. |
| Prrm4:rrm4- mKate2_ HA_G418R | pUMa4698 | genitR (G418 resistance) | Plasmid vector for generating *rrm4* C-terminal fused with mKate2_HA, construct under control of the endogenous promotor of *rrm4*. |
| Prrm4:Rrm4_mR1_ mKate2_HA_ G418R | plasmid: pUMa4569 | genitR (G418 resistance) | Plasmid vector for generating a mutant version of *rrm4* C-terminal fused with mKate2_HA. mutation in RRM1: TVEF (116-119 aa) to AAA, construct under control of the endogenous promotor of *rrm4*. |
| Prrm4:Rrm4- mR123- mKate_HA_G418R | plasmid: pUMa4571 | genitR (G418 resistance) | Plasmid vector for generating a triple mutant version of *rrm4* C-terminal fused with mKate2_HA. Rrm4 C-terminally fused to mKate2-HA; mutation in RRM1: TVEF (116-119 aa) to AAA; in RRM2: VLEF (201-205 aa) to AAAA, in RRM3: FVSF (362-368 aa) to AAAA, endogenous promotor of *rrm4.* |
| Prrm4:rrm4_mR3_R fp_ HygR | pUMa1703 | HygR (Hygromycin resistance) | Plasmid vector for generating a mutant version of *rrm4* C-terminal fused with Rfp. Mutation in RRM3: FVSF (362-368 aa) to AAAA, construct under control of the endogenous promotor of *rrm4*. |

**Table S3: DNA oligonucleotides used in this study**

| Designation | Nucleotide sequence (5' → 3') | Experiment/ Validation of: |
|---|---|---|
| MF225 | AATTGACCGCACCGTGGG | pUMa636/ pUL192, pUMa1703 |
| MF502 | ACGACGTTGTAAAACGACGGCCAG | pUMa4571, pUMa4569 pUL172, pUL173, pUMa4698 |
| MF233 | TTGAGCTGGCCAACGAGC | pUMa4571 |
| SL56 | CCCACATCCGCTCTAACC | pUMa4571, pUMa4569 |
| UM805 | TTCCGAGGTGCACTTGAAG | pUMa4569,pUL192, pUL193, pUMa4698 |
| CD503 | CATGCCATGGGAGCTAGCGCGGCCGCCAGCTCCGGTCTGCCTCTCC | pUL193 |
| EF281 | CTGGGCGTCTTGTTCAGAGT | RT-PCR: rrm4 |
| EF282 | GCCACCATGACTGACAAGGA | RT-PCR: rrm4 |
| EF283 | GGCCTTCTACGTCGCTATCC | RT-PCR: actin |
| EF284 | TCGAGACGCAGAATCGAGTG | RT-PCR: actin |
| L3-App | /rApp/AGATCGGAAGAGCGGTTCAG/ddC/ | iCLIP2 |
| RT_iCLIP2 | GGATCCTGAACCGCT | iCLIP2 |
| L01_iCLIP2.0 | /5Phos/NNNNATCACGNNNNNAGATCGGAAGAGCGTCGTG/3ddC/ | iCLIP2 |
| L02_iCLIP2.0 | /5Phos/NNNNCGATGTNNNNNAGATCGGAAGAGCGTCGTG/3ddC/ | iCLIP2 |
| L03_iCLIP2.0 | /5Phos/NNNNTTAGGCNNNNNAGATCGGAAGAGCGTCGTG/3ddC/ | iCLIP2 |
| L04_iCLIP2.0 | /5Phos/NNNNTGACCANNNNNAGATCGGAAGAGCGTCGTG/3ddC/ | iCLIP2 |
| L05_iCLIP2.0 | /5Phos/NNNNACAGTGNNNNNAGATCGGAAGAGCGTCGTG/3ddC/ | iCLIP2 |
| L06_iCLIP2.0 | /5Phos/NNNNGCCAATNNNNNAGATCGGAAGAGCGTCGTG/3ddC/ | iCLIP2 |
| L07_iCLIP2.0 | /5Phos/NNNNCAGATCNNNNNAGATCGGAAGAGCGTCGTG/3ddC/ | iCLIP2 |
| L08_iCLIP2.0 | /5Phos/NNNNACTTGANNNNNAGATCGGAAGAGCGTCGTG/3ddC/ | iCLIP2 |
| L09_iCLIP2.0 | /5Phos/NNNNGATCAGNNNNNAGATCGGAAGAGCGTCGTG/3ddC/ | iCLIP2 |
| L10_iCLIP2.0 | /5Phos/NNNNTAGCTTNNNNNAGATCGGAAGAGCGTCGTG/3ddC/ | iCLIP2 |
| L11_iCLIP2.0 | /5Phos/NNNNATGAGCNNNNNAGATCGGAAGAGCGTCGTG/3ddC/ | iCLIP2 |
| L12_iCLIP2.0 | /5Phos/NNNNCTTGTANNNNNAGATCGGAAGAGCGTCGTG/3ddC/ | iCLIP2 |
| L13_iCLIP2.0 | /5Phos/NNNNAGTCAANNNNNAGATCGGAAGAGCGTCGTG/3ddC/ | iCLIP2 |

| | | |
|---|---|---|
| L14_iCLIP2.0 | /5Phos/NNNNAGTTCCNNNNNNAGATCGGAAGAGCGTCGTG/3ddC/ | iCLIP2 |
| L15_iCLIP2.0 | /5Phos/NNNNATGTCANNNNNNAGATCGGAAGAGCGTCGTG/3ddC/ | iCLIP2 |
| L16_iCLIP2.0 | /5Phos/NNNNCCGTCCNNNNNNAGATCGGAAGAGCGTCGTG/3ddC/ | iCLIP2 |
| L17_iCLIP2.0 | /5Phos/NNNNCAACTANNNNNNAGATCGGAAGAGCGTCGTG/3ddC/ | iCLIP2 |
| L18_iCLIP2.0 | /5Phos/NNNNGTCCGCNNNNNNAGATCGGAAGAGCGTCGTG/3ddC/ | iCLIP2 |

### Table S4. Identifiers of selected RRM12-BS containing target transcripts

| Name | UMAG number |
|---|---|
| Pdc1/Bmh1 | UMAG_01366 |
| Cda4 | UMAG_01143 |
| Cdc42 | UMAG_00295 |
| Rac1 | UMAG_00774 |
| Rho1 | UMAG_05734 |
| Cdc3 | UMAG_10503 |
| Cdc10 | UMAG_10644 |
| Cdc11 | UMAG_03449 |
| Cdcd12 | UMAG_03599 |
| Chs8/Mcs1 | UMAG_03204 |
| Rax2 | UMAG_05194 |
| Atp16 | UMAG_01103 |
| Mdh1 | UMAG_10276 |
| Tom40 | UMAG_00614 |
| F-actin | UMAG_11177 |
| Chs1 | UMAG_10718 |
| Chs2 | UMAG_04290 |
| Chs3 | UMAG_10120 |
| Chs4 | UMAG_10117 |
| Chs5 | UMAG_10277 |
| Chs6 | UMAG_10367 |
| Chs7 | UMAG_05480 |
| Chs8/ Mcs1 | UMAG_03204 |
| 1,3-β-glucan synthase | UMAG_01639 |
| Cts1 | UMAG_10419 |
